# Description of *Gopheromyces tardescens*, gen. nov., sp.nov., *Gigasporangiomyces pilosus,* gen. nov., sp.nov., *Kelyphomyces adhaerens,* gen. nov., sp. nov., proposal of *Testudinimycetales* ord. nov. and *Testudinimycetaceae* fam. nov., and emended description of the order *Neocallimastigales*

**DOI:** 10.1101/2025.11.20.689491

**Authors:** Alexandria Morris, Taylor Mills, Samuel L. Miller, Carrie J. Pratt, Yan Wang, Mostafa S. Elshahed, Noha H. Youssef, Julia Vinzelj

## Abstract

Anaerobic gut fungi (AGF, *Neocallimastigomycota*) represent a phylum of zoospore-producing fungi inhabiting the gastrointestinal tracts of herbivores. Twenty mammalian-affiliated genera (M-AGF) and two tortoise-affiliated genera (T-AGF) have been described so far. Here, we report on three additional novel T-AGF isolates obtained from Texas and sulcata tortoises. Phylogenetic analysis using the D1-D2 regions of the large ribosomal RNA subunit (D1-D2 LSU), RNA polymerase II large subunit (RPB1), internal transcribed spacer-1 region (ITS1), and transcriptomics-enabled phylogenomic analysis clustered these strains into three distinct, deep-branching clades, closely related to previously described T-AGF genus *Testudinimyces*. All isolates displayed filamentous rhizoidal growth patterns and produced monoflagellated zoospores. Unique morphological characteristics included the production of elongated, thick, nucleated structures in GX isolates, the formation of thin hair-like projections on sporangial walls in SR isolates, and irregularly shaped sporangia in TM isolates. All strains grew optimally at 32-35 °C and showed distinct substrate utilization capacity (e.g., growth on pectin, chitin, galactose). LSU analyses revealed GX isolates as the first cultured representatives of tortoise-affiliated but previously uncultured lineage NY56, while SR and TM strains have not been encountered in prior culture-independent AGF surveys. We propose to accommodate these isolates in three new genera and species – *Gopheromyces tardescens* (GXA2), *Gigasporangiomyces pilosus* (SR0.6), and *Kelyphomyces adhaerens* (TM0.3). Further, based on the ecological, physiological, and phylogenetic distinctions between T-AGF and M-AGF, we propose to establish a new family (*Testudinimycetaceae*) to accommodate the genera *Testudinimyces, Gopheromyces*, *Gigasporangiomyces,* and *Kelyphomyces*, within a new order (*Testudinimycetales*), and amend the description of *Neocallimastigales* to circumscribe M-AGF genera only.

## Introduction

Anaerobic gut fungi (AGF) reside in the gut of herbivorous animals and are the only fungi known to date that are strict anaerobes.^1,2^ They belong to an early-branching phylum (*Neocallimastigomycota*), currently encompassing one class (*Neocallimastigomycetes*), one order (*Neocallimastigales*), four families, 22 genera, and 38 species.^3–6^ AGF are nutritional symbionts, aiding in the digestion of plant biomass.^1,2^ They were first discovered in domesticated mammalian hosts (e.g. sheep, cattle, horses)^7–10^ and subsequent efforts demonstrated their wide distribution in mammalian foregut and hindgut fermenters.^11–15^

In addition to mammals, some reports have documented the occurrence of AGF in non-mammalian herbivorous hosts, e.g. lizards (green iguana)^16^, birds (ostriches)^6,17^, and tortoises.^18^ Two recently described AGF genera (*Testudinimyces* and *Astrotestudinimyces*) obtained from tortoise fecal samples represent the first AGF isolates from a non-mammalian host.^4^ These genera displayed an ecological preference to tortoises, representing the predominant (or sole) AGF taxa in tortoise fecal samples examined, while being absent (or sporadically present in extremely rare relative abundance) in mammalian feces.^15,18^ Given the limited number of tortoise fecal samples examined so-far (n = 11)^18^, we hypothesized that more novel tortoise-affiliated AGF (T-AGF) lineages would be encountered when examining additional tortoise fecal samples.

Here, we report on the isolation and characterization of three novel genera from Texas and sulcata tortoise fecal samples, expanding the total number of T-AGF genera from two to five. We propose the accommodation of four of the five T-AGF into one new family (*Testudinimycetaceae*). A new order (*Testudinimycetales*) is also proposed, and we amend the description of the order *Neocallimastigales* to solely circumscribe M-AGF genera.

## Material and Methods

### Samples

Fresh fecal samples were obtained from a Texas tortoise (*Gopherus berlandieri*) at the Oklahoma City Zoo (Oklahoma City, Oklahoma, USA) in November 2020 and November 2024, from a sulcata tortoise (*Centrochelys sulcata*) at the Tanganyika Wildlife Park (Goddard, Kansas, USA) in August 2024, and from another sulcata tortoise at the Amarillo Zoo (Amarillo, Texas, USA) in February 2025. Samples were collected in sterile 50 mL Falcon tubes, frozen on site and transported to the laboratory, where they were stored at -20°C until used for isolation.

### Isolation

Multiple isolation efforts were conducted between June 2022 and April 2025 (Table S1). Enrichments were set up in an anaerobic chamber (Coy Laboratories, Grass Lake, MI, USA) on rumen-fluid cellobiose medium (RFC) supplemented with antibiotics (50 μg/mL chloramphenicol, 50 μg/mL penicillin G sodium salt, 20 μg/mL streptomycin sulfate) and switchgrass.^19^ Enrichments were incubated stationary in the dark at 30 °C and monitored daily for signs of fungal growth (clumping and/or floating of the switchgrass, biomass formation, and sticking to glass, production of gas bubbles). Once growth was observed, roll tubes (RFC supplemented with antibiotics and 2% agar)^20^ were prepared from the enrichment tubes and incubated again until colony growth was observed. Single colonies were then picked in an anaerobic chamber and transferred to RFC medium supplemented with antibiotics. Roll tubing and colony-picking were repeated twice to ensure purity.

### Morphological characterization

Morphological features of all isolates were examined by phase contrast microscopy using a BX51 (Olympus, Center Valley, PA, USA) instrument equipped with an MU503-GS AmScope digital camera; confocal microscopy, using Zeiss LSM 980 Airyscan 2 confocal laser scanning microscope; and scanning electron microscopy using an FEI Quanta 600 field-emission gun environmental scanning electron microscope with a Bruker EDS X-ray microanalysis system and an HKL EBSD system. To examine thallus structure and nuclear localization, samples were stained with 4,6′-diamidino-2-phenylindole (DAPI; 10 µg/mL) and visualized using phase-contrast and confocal laser-scanning microscopy. For electron microscopy, samples were fixed and dried as previously described.^21^ Briefly, the medium of 4-day-old cultures (grown in Balch tubes in 10 mL RFC medium) was carefully removed with Pasteur pipettes before biomass was transferred into 2 mL plastic tubes. Biomass was fixed for 2 hours at room temperature (∼23 °C) in 2% glutaraldehyde dissolved in 0.1 M cacodylate buffer. After three 15-minute washing steps in a buffered wash, cells were fixed in 1% OsO_4_ for 1 hour at room temperature. Washing with buffered wash was repeated before sequential dehydration in ethanol (50%, 70%, 80%, 90%, 95%, 3x 100%) for 15 minutes each. The samples were further dried twice in hexamethyldisilane for five minutes each and left to dry overnight. At no point during the process were the samples centrifuged to prevent loss of delicate structures. Samples were then mounted on stubs, Au-Pd coated, and imaged. Sizes of various microscopic structures were measured using Fiji software.^22^

### Substrate utilization and temperature preferences

To assess substrate utilization patterns, cultures were grown in Balch tubes in rumen fluid media containing various carbon sources with a total volume of 10 mL (7 mL media and 3 mL fungal culture) and followed for four consecutive subcultures. Media were supplemented with antibiotics (50 μg/mL chloramphenicol, 50 μg/mL penicillin, and 20 μg/mL streptomycin), and the respective carbon source (0.5% w/v). Three replicates were tested per substrate and strain, along with uninoculated controls. Tubes were incubated at 32 °C (strain TM0.3), or 35 °C (strains GXA2 and SR0.6) for four (strains SR0.6 and TM0.3), or seven (strain GXA2) days before growth was assessed and the tubes subcultured. Growth was evaluated by visual inspection (biomass production, biomass sticking to glass, turbidity indicating bacterial contamination, redness of the media indicating oxygenation) and gas pressure measurements (MediaGauge, SSI Technologies, Wisconsin, USA). Gas pressure readings were conducted at 35 °C to eliminate any influence temperature changes might have on the pressure reading. Gas pressure measurements and visual growth evaluation were condensed into a qualitative growth scale of – (no growth), + (very slight growth), ++ (slight growth), +++ (good growth), and ++++ (very good growth), given that visually well-growing cultures did not produce noticeable gas pressure on some substrates (Table S2).

To ascertain the optimal growth temperature, isolates were grown in RFC medium containing a preferred substrate (lactose for GXA2, cellobiose for SR 0.6 and TM0.3) as the sole carbon source. Cultures were incubated at six different temperatures (22 °C, 28 °C, 32 °C, 35 °C, 37 °C, and 39 °C), and growth was evaluated as described above. All tubes were acclimatized to 35 °C before gas pressure readings were recorded. Per isolate, three replicates and one uninoculated control were investigated at each temperature for six consecutive subcultures.

The data for growth curves was obtained through daily evaluation of three tubes per strain grown on the preferred substrate at the optimal temperature.

### DNA extraction and phylogenetic analysis

DNA was extracted from cultures grown in RFC medium until peak biomass production (strain GXA2, seven days; strain SR0.6, four days; strain TM0.3, five days) using the DNeasy PowerPlant Pro Kit (Qiagen Corp., Germantown, MD, USA) following the manufacturer’s instructions. For initial identification, the D1-D2 region of the 28S rRNA gene (D1-D2 LSU) was amplified using primers NL1 (5′-GCATATCAATAAGCGGAGGAAAAG-3′)^23^ and NL4 (5′-GGTCCGTGTTTCAAGACGG-3′)^23^ in a 25 µL PCR reaction using the DreamTaq PCR Master Mix (ThermoFisher Scientific, Waltham, MA, USA) according to the manufacturer’s instructions. The PCR program consisted of an initial denaturation of 5 min at 95 °C followed by 40 cycles of denaturation at 95 °C for 30 seconds, annealing at 52 for 30 sec, and elongation at 72 for 1 minute. The final elongation step was run at 72 for 10 minutes. PCR products were cleaned using the PureLink™ PCR Purification Kit (Invitrogen, Carlsbad, CA, USA) and sent for Sanger sequencing at the Oklahoma State University (OSU) DNA Protein Core Facility (Stillwater, OK, USA).

For a more thorough phylogenetic analysis, the extracted DNA was also used as a template to amplify the region encompassing ITS-1, 5.8S rRNA, ITS-2, and the D1/D2 domains of the 28S (LSU) rRNA gene (approximately 1.3 kb) using the primers ITS5F (5’-GGAAGTAAAAGTCGTAACAAGG-3’; clade GX)^24^ or ITS1 (5’-TCCGTAGGTGAACCTGCGG-3’; clades SR and TM)^24^ and NL4 using the same protocol as described above. Cleaned PCR products were cloned into a TOPO-XL2 cloning vector (Life Technologies®, Carlsbad, CA, USA) following the manufacturer’s instructions. Per strain, five to 12 clones were Sanger-sequenced at the OSU DNA Protein Core Facility (Stillwater, OK, USA; strain GX1) or sequenced by Plasmidsaurus (strains SR0.6 and TM0.3, Oxford nanopore sequencing with v14 chemistry on an R10.4.1 flow cell, USA). ITS1 and D1-D2 28S rRNA sequences were extracted from the sequenced clones using MEGA.^25^ Recently obtained genomic sequences of the type strains (unpublished data) were also mined for rRNA loci using local Blastn, and the regions corresponding to ITS1 and D1-D2 28S rRNA sequences were extracted using BEDTools.^26^ Sequences (clones or genomic) were aligned to reference ITS1 and D1-D2 LSU sequences (AFN database v2, www.anaerobicfungi.org/databases) using MAFFT^27^, and *Chytriomyces* sp. WB235A isolate AFTOL-ID 1536 as an outgroup. For protein trees, sequences of the RNA polymerase II large subunit gene (RPB1) were bioinformatically extracted from the transcriptomes of the isolates (obtained as described below). Protein sequences were aligned to reference sequences using MAFFT.^27^

IQ-TREE^28^ was used to predict the best substitution model and to generate maximum-likelihood trees under the predicted best model. Options ‘–alrt 1000’, ‘-abayes’, and ‘–bb 1000’ were added to the command line to perform the Shimodaira–Hasegawa approximate-likelihood ratio test (SH-aLRT), approximate Bayes tests, and ultrafast bootstrap (UFB), respectively. IQ-TREE analysis resulted in the generation of phylogenetic trees with three support values (SH-aLRT, aBayes, and UFB) for each branch.

### Transcriptomic sequencing, AAI calculation, phylogenomic analysis, and molecular timing

Total RNA was extracted from six 10-mL cultures grown in RFC media to late exponential/ early stationary phase at peak-biomass production (strain GXA2, seven days; strain SR0.6, four days; strain TM0.3, 5 days). Biomass was vacuum-filtered, and cells were lysed by crushing in liquid nitrogen. Total RNA was extracted using the NucleoSpin™ RNA Mini Kit (Macherey-Nagel™) according to the manufacturer’s instructions. Total RNA was sequenced on an Illumina NextSeq 2000 platform using a 2×150 bp paired-end library at the One Health Innovation Foundation lab at OSU.

Transcriptomes were quality-trimmed and *de-novo* assembled from RNA-seq reads using Trinity (version 2.6.6)^29,30^ with default parameters. CD-HIT^31^ was used to cluster redundant transcripts with an identity parameter of 95 % (–c 0.95). Peptide and coding sequences were predicted using TransDecoder (version 5.0.2; https://github.com/TransDecoder/TransDecoder) with a minimum peptide length of 100 amino acids. The predicted peptides were used to extract the single-copy protein RPB1 for phylogenetic assignment (see above), average amino acid identity (AAI), as well as for phylogenomic analysis.

For AAI calculations, we included predicted peptides from other previously obtained AGF transcriptomes (*n* = 60).^15,18,32^ AAI values were calculated for all possible pairs in the dataset using the *aai.rb* script available as part of the Enveomics collection (https://github.com/lmrodriguezr/enveomics).^33^

For phylogenomics and molecular dating analysis of evolutionary divergence, we included the transcriptomics data of newly obtained isolates and 60 available AGF transcriptomes^15,18,32^, in addition to 5 *Chytridiomycota* genomes (*Chytriomyces* sp. strain MP 71, *Entophlyctis helioformis* JEL805, *Gaertneriomyces semiglobifer* Barr 43, *Gonapodya prolifera* JEL478, and *Rhizoclosmatium globosum* JEL800). Profile hidden Markov models (HMMs) of the 758 phylogenomic markers for Kingdom Fungi in the ‘fungi_odb10’ dataset^34^ were used for analysis as previously described.^15,18,31^ The HMMs and the PHYling pipeline^35^ (https://doi.org/10.5281/zenodo.1257001) were used to identify homologues in all AGF transcriptomes (n = 63), as well as the five *Chytridiomycota* genomes using HMMER3 (http://hmmer.org/). Markers with conserved homologs in all datasets were aligned and concatenated. The refined alignment was grouped using PartitionFinder (v 2.1.1)^36^ to assign each partition with an independent substitution model. BEAUti (v 1.10.4)^37^ was used for Bayesian and molecular dating analyses using two calibration priors: a direct fossil record of *Chytridiomycota* from the Rhynie Chert (407 Mya) and the emergence time of *Chytridiomycota* (573 to 770 Mya as 95% HPD). Three independent runs (30 million generations each) were performed with a default burn-in (10%). The Birth-Death incomplete sampling tree model was used for interspecies relationship analyses. Unlinked strict clock models were used for each partition independently. Tracer (v1.7.1)^38^ was then used to confirm that a sufficiently effective sample size (ESS > 200) was obtained. Finally, the maximum clade credibility (MCC) tree was compiled using TreeAnnotator (v1.10.4).^37^

### Data and culture accession

Sequences generated in this study are deposited in GenBank under the BioProject accession number PRJNA1345044. Cultures are available at Oklahoma State University, Department of Microbiology and Molecular Genetics culture collection (Stillwater, OK, USA), where they are kept in active culture (30 °C, Balch tubes with RFC medium, subcultured weekly) as well as preserved at -80 °C (following protocol LN described in Vinzelj et al., 2022).^39^

## Results

### Isolation

Enrichments from tortoise fecal samples yielded six isolates (Table 1, Table S1): three from an OKC Zoo Texas tortoise (GX1, GX2, GXA2), two from a Tanganyika Wildlife Park sulcata tortoise (SR0.1 and SR0.6), and one from an Amarillo Zoo sulcata tortoise (TM0.3) (Table S1). Isolation efforts from the Texas tortoise were initiated after a culture-independent survey of the fecal sample indicated a near predominance (98.31%) of a potentially novel genus-level clade of AGF (Table S1 in Pratt *et al.*, 2024).^18^ Three separate isolates were obtained in June 2022 (isolate GX1), November 2024 (isolate GX2), and January 2025 (isolate GXA2) (Table S1). Strain GX1 lost viability three months after isolation. Strains GX2 and GXA2 remained viable, albeit slow growing, reaching mid-log phase (near-peak biomass production) six to eight days after subculturing (Figure S1). Biomass production slows down drastically when grown continuously on a single substrate for one to two months (with weekly transfers into new media). Regular alternation of the carbon source, however, appears to aid in retaining the strains’ viability. Isolate GXA2 was chosen as the type strain for further characterization.

**Table 1:**
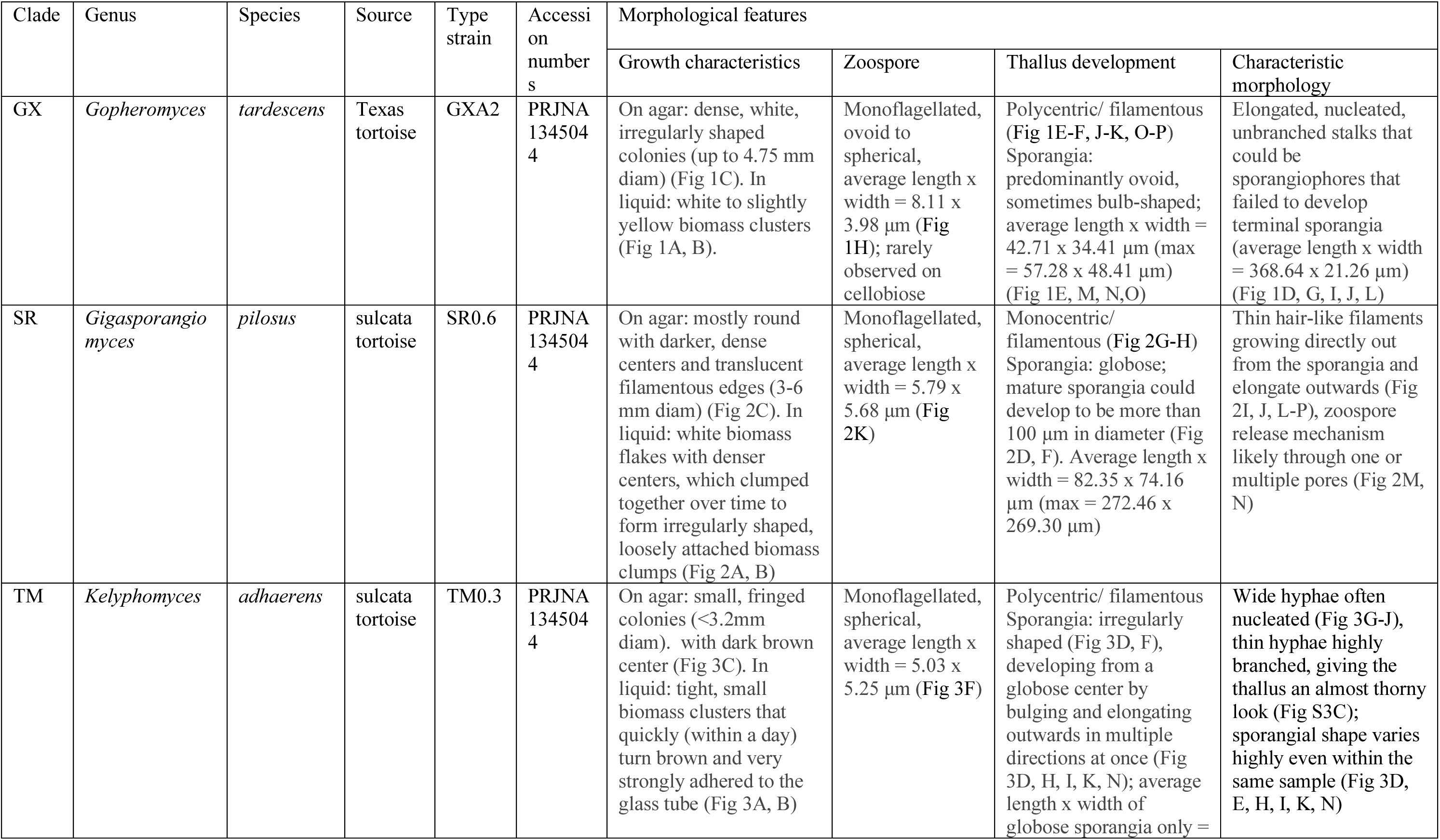
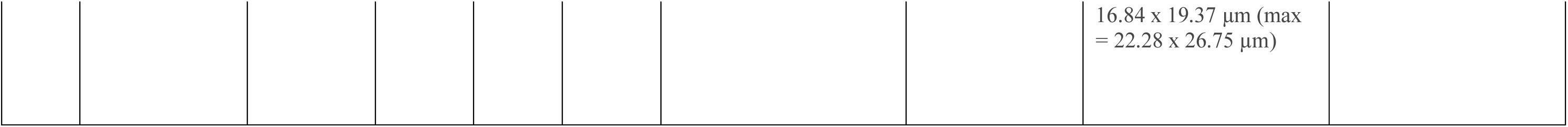
Overview on the novel clades of *Neocallimastigomycota* described in this study.

Strains SR0.1 and SR0.6 were isolated in March 2025. The effort was initiated after a culture-independent survey of the fecal sample indicated near predominance of potentially novel genus-level clades of AGF (Table S2). The growth rate of both strains was faster than that of the GX isolates, reaching near-peak biomass production after two to four days of incubation (Figure S1). Both strains have remained viable on RFC media since their isolation with weekly transfers at 30 °C. Isolate SR0.6 was chosen as the type strain for further characterization.

Strain TM0.3 (type strain) was obtained in April 2025, with the effort also initiated by results obtained from a culture-independent survey (Table S2). The growth rate of strain TM0.3 was comparable to the SR strains, reaching near-peak biomass production after three to five days of incubation (Figure S1), and is similarly still active on RFC media with weekly transfers at 30 °C.

#### Morphological characterization

##### Macroscopic features

In liquid RFC media, isolate GXA2 produced white to slightly yellow biomass clusters (Figure 1A) that adhered to the glass tubes but detached upon shaking. In rumen-fluid media containing lactose as sole carbon source, GXA2 formed thin biomass films that stuck tightly together and to the glass (Figure 1B). On RFC agar roll tubes, the strain produced irregularly shaped, white colonies that started out small and grew over time (up to 4.76 mm in diameter). The dense center darkened with age, while the fringed edges remained white to yellowish.

**Figure 1:**
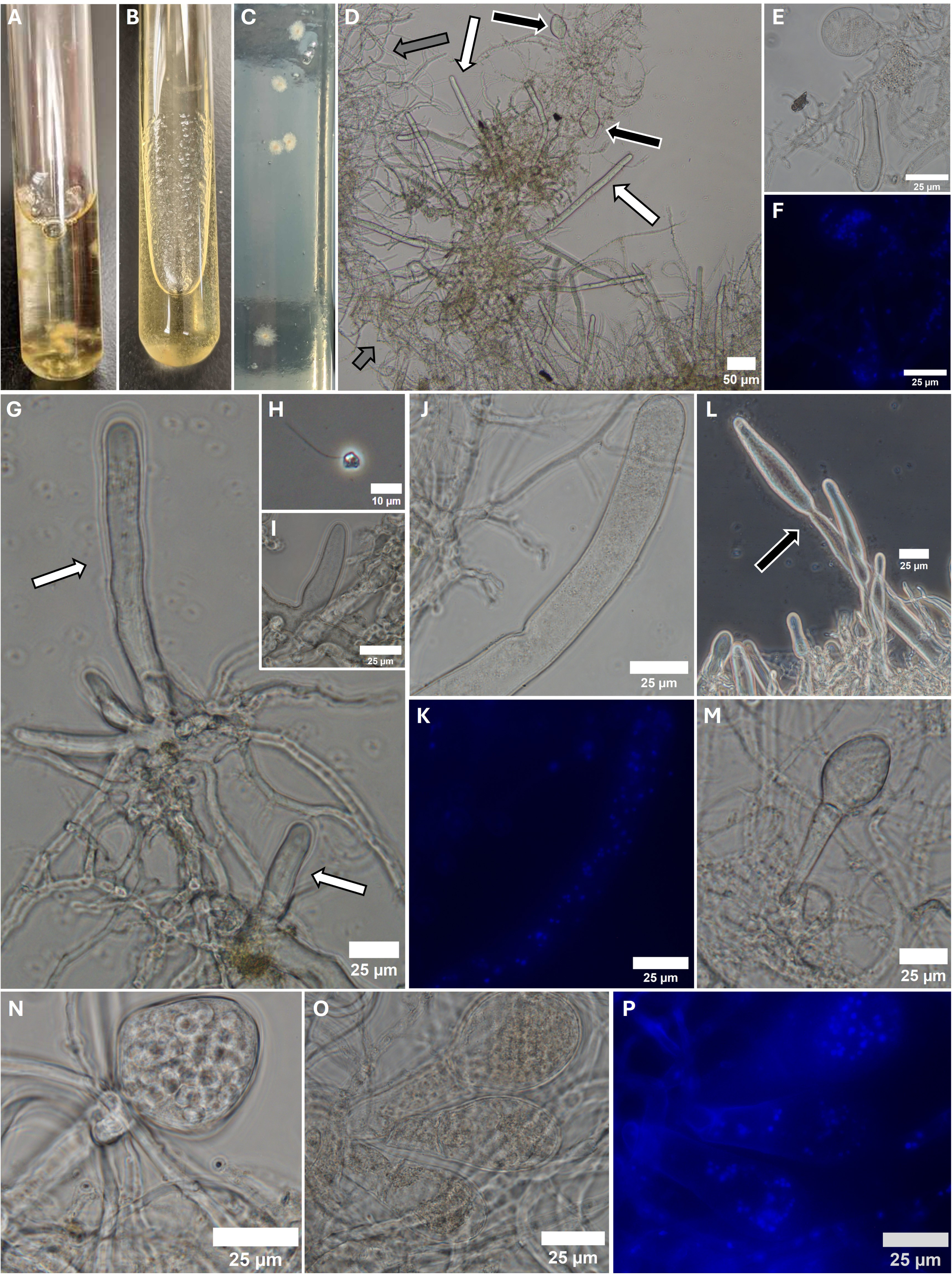
Morphology of strain GXA2 (Clade GX, *Gopheromyces tardescens)*. **(A)** Strain GXA2 grown for five days in rumen fluid cellobiose (RFC) medium. **(B)** Strain GXA2 in rumen fluid medium with lactose instead of cellobiose. **(C)** Morphology on RFC agar rolltubes. **(D-P)** Morphological features of GXA2 grown in rumen fluid media containing either cellobiose **(J, K, O, P)** or pectin as the sole carbon source. **(D)** The features of GXA2: filaments with wide and narrow hyphae (grey arrow), long unbranching stalks (likely sporangiophores that failed to produce terminal sporangia; white arrows), and sporangia (black arrows). **(E/F, J/K, O/P)** DAPI staining showing nucleated sporangia, hyphae, and sporangiophores. **(E, I, G)** Development of sporangiophores (white arrows) out of hyphae. **(H)** Monoflagellated zoospore. **(L)** Sporangiophores pinching (black arrow) and swelling of the tip. **(M)** Typical developing sporangium. **(N)** Zoospore-filled sporangium. Scale bars indicate 50 µm **(D)**, 25 µm **(E-G, I-P)**, or 10 µm **(H)**.

In liquid RFC media, isolate SR0.6 produced white biomass flakes with denser centers, which clumped together over time to form irregularly shaped, loosely attached biomass clumps that detached when shaken (Figure 2A, B). On RFC agar roll tubes, the strain formed round colonies (3.2 to 4.8 mm in diameter), with white to yellowish filamentous edges that had a translucent sheen. The small, dense centers started out the same color as the edge and darkened over time (Figure 2C).

**Figure 2.**
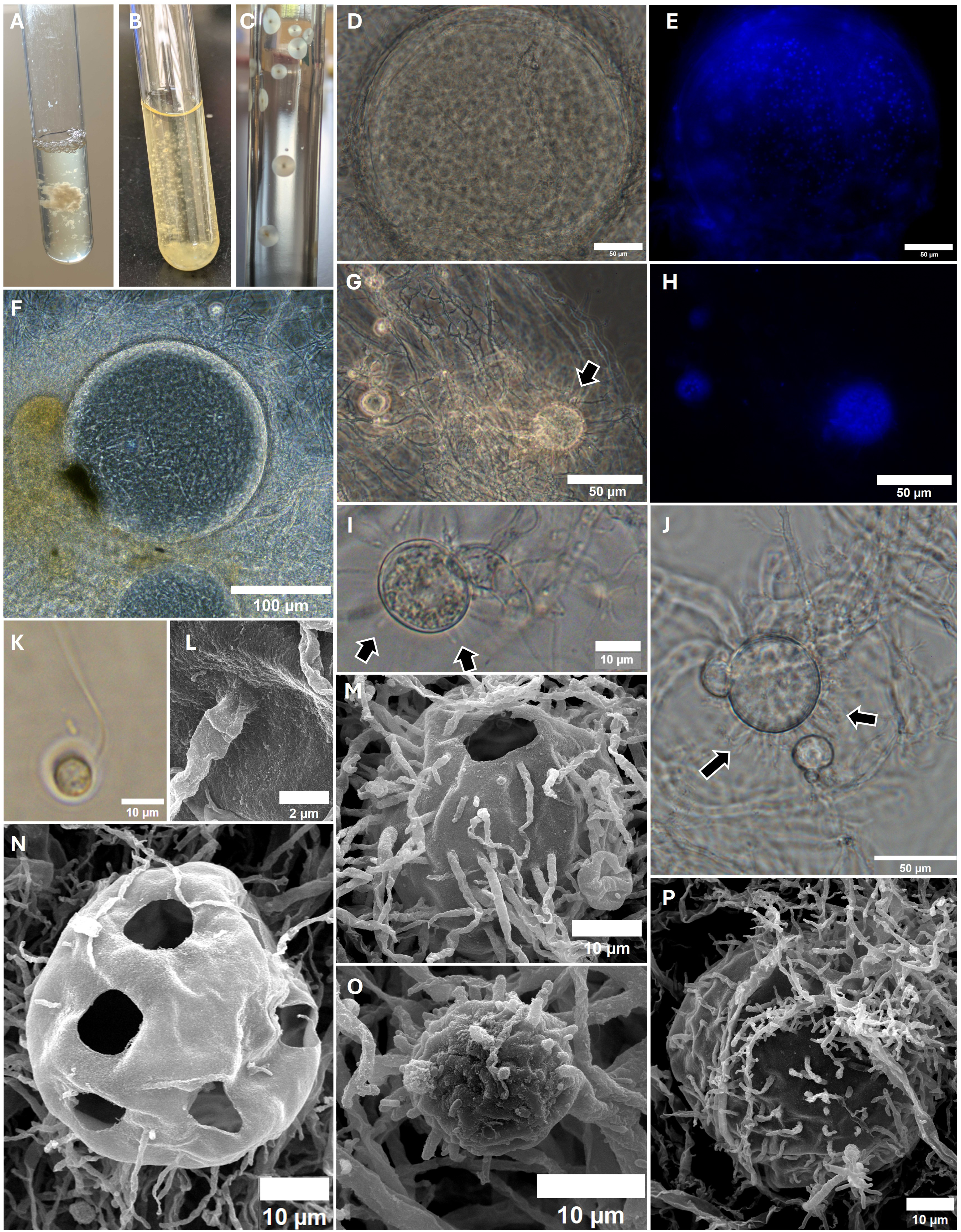
Morphology of isolate SR0.6 (Clade SR, *Gigasporangiomyces pilosus)*. **(A, B)** Strain SR0.6 grown for four days in rumen fluid cellobiose (RFC) medium. **(A)** before shaking, **(B)** after shaking. **(C)** Morphology on RFC agar rolltubes. **(D-K)** Phase contrast microscopy of strain SR0.6. **(D/E, G/H)** DAPI staining showing the nuclei present in sporangia but absent in hyphae. **(D-F)** Giant sporangia (>100 µm) characteristic for this strain. **(G, I, J)** Developing sporangia showing thin, hair-like structures growing out of the sporangial body (black arrows). **(K)** Monoflagellated zoospore. **(L-P)** Scanning electron microscopy of strain SR0.6 grown in RFC for four days. The thin, hair-like structures growing out of the sporangia are visible here as well. **(L)** Close-up of the thin, hair-like filaments growing out of the sporangial wall. **(M, N)** Pores found on sporangia that could stem from zoospore release. Scale bars: **(F)** 100 µm, **(D-E, G-H, J)** 50 µm, **(I, K, M-P)** 10 µm, **(L)** 2 µm.

In liquid RFC media, strain TM0.3 produced tight, small biomass clusters that quickly darkened and adhered very strongly to the glass tube (Figure 3A, B). Rough shaking was necessary to detach and break enough biomass for the transfer of strain TM0.3, and complete removal of biomass from the tubes was only possible by physically scraping the biomass from the tubes. (Figure 3B). On RFC agar roll tubes, the strain formed small colonies (<3.2 mm in diameter) with a dense center and irregularly growing filaments, giving them a fringed appearance. The colonies started out white to yellowish in color, and over time the center of the colonies turned brown with a white/yellowish denser perimeter remaining and thin filaments reaching out in all directions (Figure 3C).

**Figure 3.**
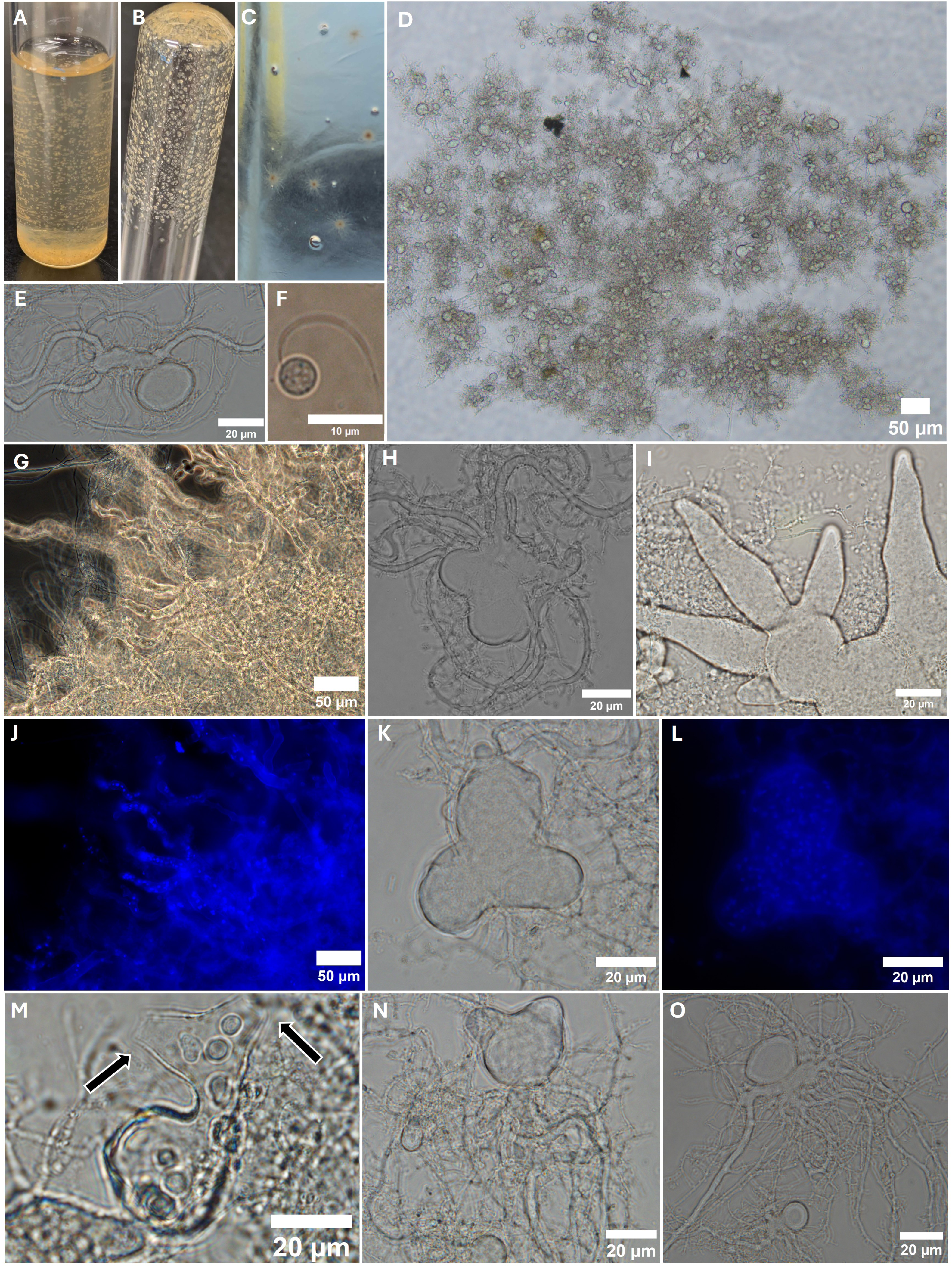
Morphology of isolate TM0.3 (Clade TM, *Kelyphomyces adhaerens*). **(A,B)** Strain TM0.3 grown for four days in liquid rumen fluid cellobiose (RFC) medium. **(C)** Morphology on RFC agar rolltubes. **(D-O)** Phase contrast microscopy of strain TM0.3 grown in liquid RFC medium **(D, I, N)** or rumen fluid medium containing inulin as sole carbon source **(E, F, G, H, J, K, L, M, O)**. **(G/J, K/L)** DAPI staining showing nucleated sporangia and hyphae. **(D, E, H, I, K, M, N)** Examples of the various sporangial shapes observed in this strain. Growth starts out globose **(E, O)** and grows out from there **(N, K, H, I)**, taking different shapes even within the same sample. **(F)** Monoflagellated zoospore **(M)** Zoospores in half-empty sporangia with ruptured sporangial walls (black arrows) as a potential zoospore release mechanism. Scale bars: **(D, G, J, N)** 50 µm, **(D, E, H, I, K-M, O)** 20 µm, **(F)** 10 µm.

##### Microscopic features

GX strains produced filamentous, polycentric thalli with nucleated rhizomycelia that displayed extensive branching and entanglement patterns (Figure 1D-P, S2A, C). Both narrow and wide hyphae (average width of 4.96 ± 1.95 µm, n = 251, Figure 1D) were observed. Additionally, elongated, unbranched structures with larger diameters compared to the wide hyphae (ranging from 8.15 to 36.24 µm, average of 21.26 ± 7.50 µm, n = 42) and variable lengths were observed (up to 413 µm measured, Table S2, white arrows in Figure 1D, G; Figure S2B, D, H). These structures grew from existing hyphae (sometimes in clusters of two or three, Figure 1G, I, S2H) and were always nucleated (Figure 1J-K). On rumen fluid media containing pectin as the sole carbon source, pinching was observed in some of these structures (black arrows in Figure 1L; Figure S2D), which was never observed on RFC media. The nature of these structures is unclear, though we speculate that they could represent elongated sporangiophores that failed to develop terminal sporangia. Developing sporangia were sometimes observed in rumen fluid media containing lactose, cellobiose or inulin as carbon sources, but zoospore-filled sporangia and free-swimming zoospores were more regularly observed on rumen fluid media containing pectin as the sole carbon source.

Free swimming zoospores were generally rarely observed. They were monoflagellated and mostly ovoid or globose in shape with an average length and width of 8.11 ± 1.79 µm and 3.98 ± 0.68 µm, respectively (n = 6, Figure 1H, Table S2). Sporangia, when observed, where mostly ovoid with an average length and width of 42.71 ± 7.52 µm and 34.41 ± 7.26 µm (n = 16, Table S2, Figure 1E, M, N, O, S2F, I, J), respectively, although bulb-shaped or globose sporangia were also sometimes observed (Figure 1O-P, S2E). Developed ovoid or bulb-shaped sporangia usually showed short sporangiophores that grew out of hyphae (Figure 1D, black arrows, 1E, O, S2F, G, I, J); a developed sporangium with zoospores was never observed at the end of the elongated structures mentioned above.

Isolate SR0.6 produced filamentous, monocentric thalli with mostly anucleated, extensive rhizomycelia (Figure 2D-P, Figure S3). Zoospores were monoflagellated and spherical, with average length and width of 5.79 ± 0.84 µm and 5.68 ± 0.88 µm (n = 15, Figure 2K, Table S2). Most sporangia were globose with average length and width of 45.17 ± 21.21 µm and 44.48 ± 20.98 µm (n = 25, Table S2) and often had filamentous, hair-like structures growing directly out from the sporangial wall (black arrows in Figure 2G, I, J; Figure 1L-P, S3C-F). Sporangia increased in size with age, with some reaching diameters of more than 100 µm (average of 139.99 ± 61.14 µm, n = 14; Figure 2D, F, S3B). The filamentous, hair-like structures elongated outwards during maturation of the sporangia, and sporangia are often firmly entangled in their own filaments (Figure 2D, F, P, S3B, C, F). SEM microscopy revealed one or multiple pores in the sporangial wall as a potential zoospore release mechanism (Figure 2M, N).

Strain TM0.3 produced filamentous, polycentric thalli with both narrow, highly branched, and wider, less branched but nucleated filaments (Figure 3D, E, G-J). Zoospores were monoflagellated and mostly globose to ovoid with an average length and width of 5.03 ± 0.55 µm and 5.25 ± 0.60 µm (n = 21, Table S2, Figure 3F). Sporangia were irregularly shaped (Figure 3D, E, H, I, K, M, N, S4B, D, E, F, H), starting out globose (Figure 3E) and then growing outwards in all directions and shapes (3D, H, I, K, N, S4E, H). Zoospore release was observed through a rupture in the sporangial wall (Figure 3M).

### Substrate utilization

The substrates glucose, fructose, cellobiose, cellulose, and xylan supported growth of all three isolates through four subcultures (Table 2, Figure S5, Table S4). AGF readily utilize such substrates, and they are routinely used for isolation. Lactose supported growth of GXA2 and SR06., but not TM0.3. Growth of strain GXA2 was supported on chitin, whereas strains TM0.3 and SR0.6 were unable to survive passed the first and second subculture, respectively. None of the strains survived on polygalacturonic acid or xylose. Galactose supported weak growth of SR0.6 and GXA2, but not TM0.3. Inulin supported very weak growth of GXA2 and SR0.6, but not TM0.3, and pectin kept GXA2 alive, but not TM0.3, and SR0.6 only survived two subcultures.

**Table 2:** Growth of strains GXA2, SR0.6, and TM0.3 on different substrates and temperatures. ‘-’ refers to no growth, ‘++++’ refers to very good growth.

| Substrates |  | GXA2 | SR0.6 | TM0.3 |
| --- | --- | --- | --- | --- |
| Monosaccharides | Arabinose | - | + | - |
|  | Fructose | +++ | +++ | ++ |
|  | Galactose | - | +++ | ++ |
|  | Glucose | +++ | +++ | +++ |
|  | Xylose | - | - | - |
| Disaccharides | Cellobiose | +++ | +++ | +++ |
|  | Lactose | +++ | +++ | - |
|  | Trehalose | - | - | + |
| Polysaccharides | Cellulose | +++ | +++ | +++ |
|  | Chitin | ++ | ++ | - |
|  | Inulin | ++ | ++ | - |
|  | Pectin | + | + | - |
|  | Polygalacturonic Acid | + | + | - |
|  | Xylan | ++ | ++ | ++ |
| Temperature (°C) |  | GXA2 | SR0.6 | TM0.3 |
|  | 22 | - | - | + |
|  | 28 | + | ++ | ++ |
|  | 32 | +++ | ++++ | +++ |
|  | 35 | ++++ | ++++ | ++++ |
|  | 37 | ++ | +++ | ++ |
|  | 39 | - | ++ | + |

### Temperature preferences

Strain GXA2 grew between 28 and 37 °C with optimal growth occurring at 35 °C, and no growth was observed at 39 °C (Table 2, Figure S6, Table S4). Strain SR0.6 grew between 28 and 39 °C with optimal growth at 32 and 35 °C. Growth at 39 °C dwindled within two months (data not shown). Strain TM0.3 grew between 22 and 37 °C with optimal growth at 35 °C. Growth at 39 °C was observed only in two subcultures. Interestingly, when grown at 22 °C the macroscopic morphology of strain TM0.3 changed from the quick-to-darken biomass that firmly adhered to glass, to white, thin wisps of biofilm that only loosely attached to glass.

### Phylogenetic analysis

Phylogenetic analysis using the D1-D2 region of the LSU rRNA gene placed the six isolates in three deeply-branching bootstrap-supported clades close to the two previously isolated AGF genera from tortoises (*Testudinimyces* and *Astrotestudinimyces*)^4^ (Figure 4A). The closest cultured representative to all three clades was *Testudinimyces gracilis* (93.94 ± 0.29% for clade GX, 95.07 ± 0.73% for clade SR, and 94.22 ± 0.15% for clade TM, Table 3). A lower similarity was observed when comparing these clades to *Astrotestudinimyces* (86.08 ± 0.22% for clade GX, 86 ± 0.71% for clade SR, and 85.34 ± 0.23% for clade TM). Inter-clade similarity values between all three clades ranged between 96.03 ± 0.77 % to 96.72 ± 0.29% (Table 3). Sequence similarity values to M-AGF genera were significantly lower (77.4 to 89.82%; average: 86.75 ± 1.98%).

**Figure 4.**
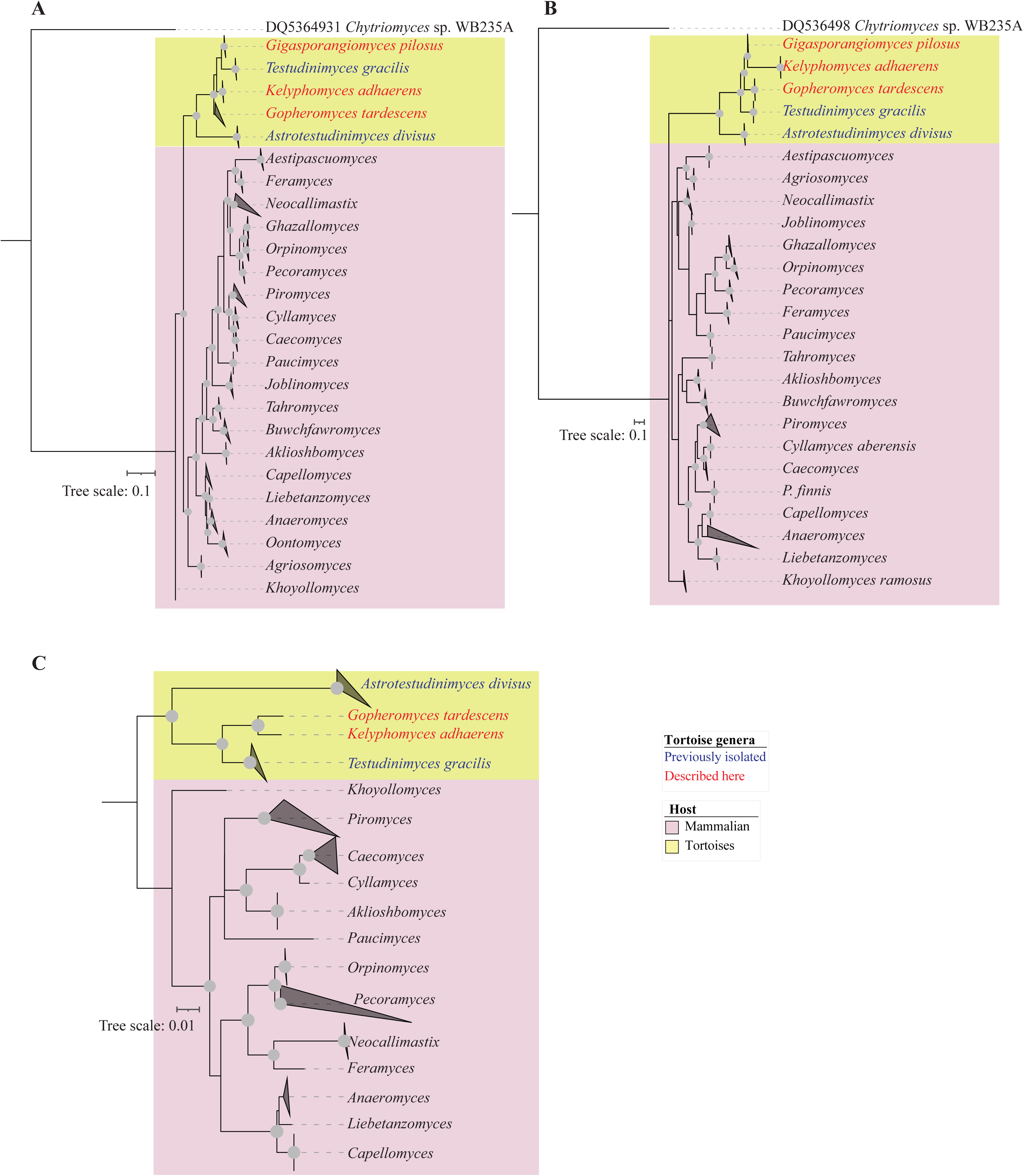
Phylogenetic trees constructed using the D1–D2 region of the LSU **(A)**, the ITS1 region **(B)**, and RPB1 **(C)** as phylogenetic markers. Alignments were created in MAFFT.^27^ Trees were constructed in IQ-TREE.^28^ using the maximum-likelihood approach*. Chytriomyces* sp. WB235A isolate AFTOL-ID 1536 was used as an outgroup in **(A)** and **(B)**. Three bootstrap support values (SH-aLRT, aBayes, and UFB) are created for each branch. Support values are shown as gray dots when all support values are above 70%. Scale bars indicate the number of substitutions per site.

**Table 3.**
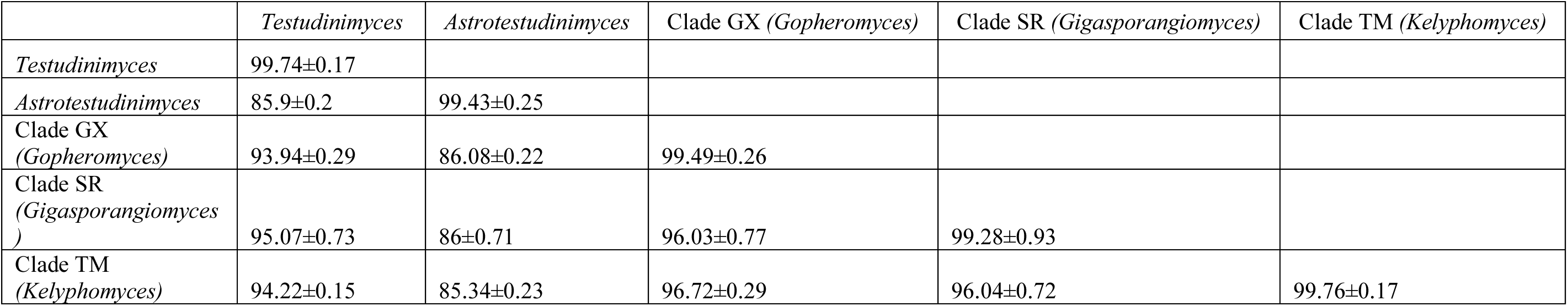
Sequence divergence in the D1-D2 region of the LSU rRNA among the five T-AGF genera.

Phylogenetic analysis using the ITS1 region also placed the six isolates in three bootstrap-supported clades, with *Testudinimyces gracilis* being their closest cultured representative (Figure 4B). Inter-clade similarity was highest for clade GX and SR, with clade TM less similar in the ITS1 region to the other two clades. Within-strain sequence divergence ranged from 0.648 ± 0.38% (clade TM) to 3.28 ± 1.96% (clade GX).

Finally, phylogenetic analysis using RPB1 placed two of the clades (GX and TM) close to *Testudinimyces* (91.872%, and 90.98%, respectively) and *Astrotestudinimyces* (92.582, and 93.472%, respectively) (Figure 4C). An RPB1 sequence was missing from SR0.6 transcriptome.

Querying D1-D2 LSU rRNA sequence data against publicly available sequences recovered from a culture-independent diversity survey of tortoise feces^18^ revealed that GX strains are the representatives of the uncultured clade NY56, previously identified as the only AGF genus in the Texas tortoise from which the strains were obtained. On the other hand, no sequences from previous culture-independent diversity surveys of tortoise feces matched those of SR or TM.

### Phylogenomics and molecular dating analysis

Transcriptomics-enabled phylogenomic analysis placed the three reported taxa in a monophyletic clade closely related to *Testudinimyces* (Figure 5), with those four genera clustering away from *Astrotestudinimyces*, further confirming the topology obtained from single-locus phylogenetic analysis (Figure 4).

**Figure 5.**
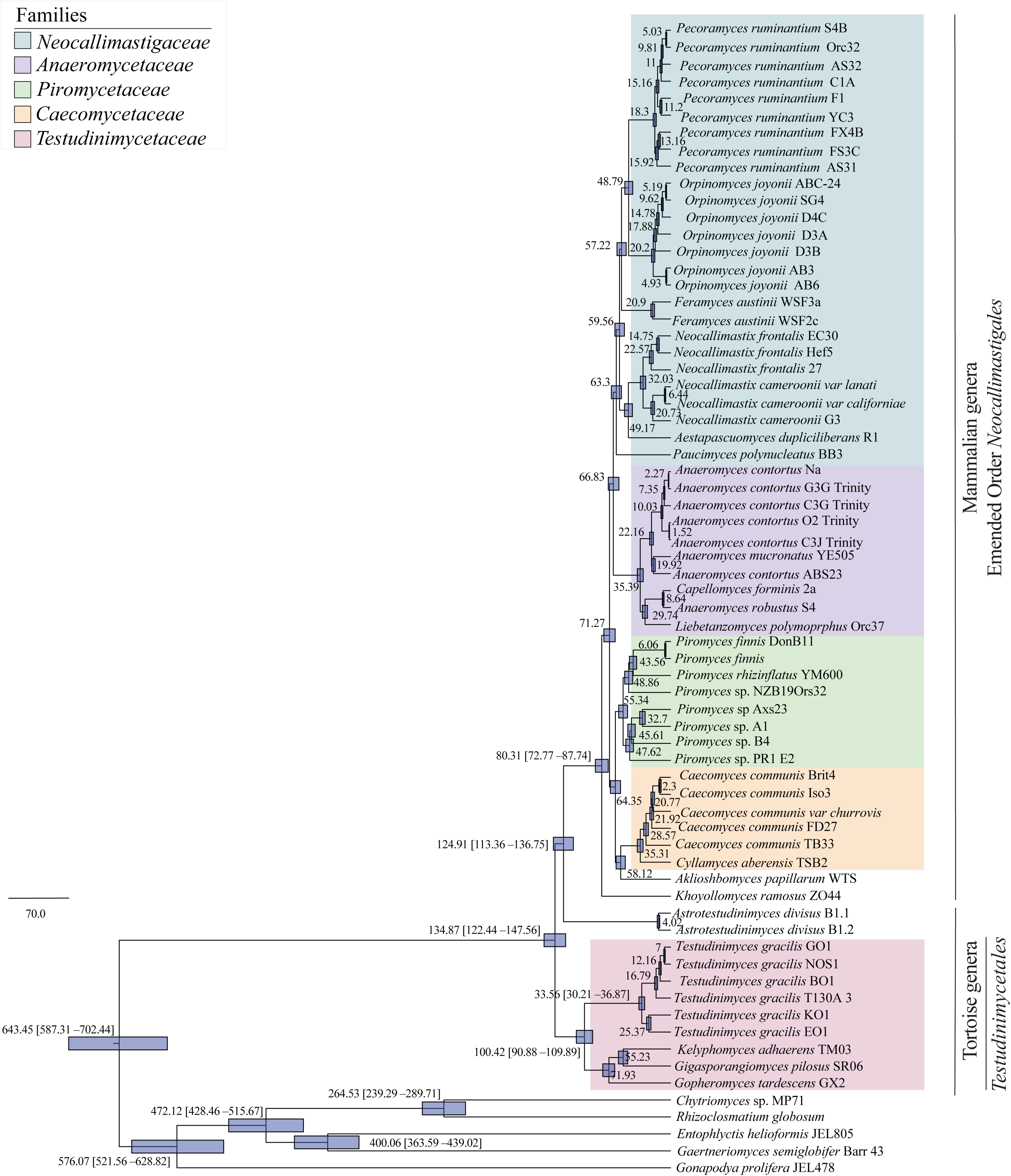
Transcriptomics-enabled phylogenomic tree of *Neocallimastigomycota* based on 758 genome-wide markers highlighting the relationships within the phylum. BEAUti (v 1.10.4)^36^ was used for Bayesian and molecular dating analyses and the maximum clade credibility (MCC) tree was compiled using TreeAnnotator (v1.10.4).^36^ The numbers at the nodes represent the estimated age in mya. Transcriptomes are collapsed at genus-level with labels at the tips being color-coded by family. Outgroups are colored in grey.

Previous molecular dating analysis with the two tortoise AGF genera *Astrotestudinimyces* and *Testudinimyces* pushed the *Neocallimastigomycota* divergence to the early Cretaceous, with the earliest divergence time estimated at 112.19 Mya (genus *Astrotestudinimyces* lineage). The addition of the three novel taxa to the AGF phylogenomic tree further suggested an earlier divergence time of AGF, to the early Cretaceous/late Jurassic (134.87 Mya; with the 95% Highest Probability Density (HPD) interval at 122.44 to 147.56 Mya). The divergence time of the genus *Astrotestudinimyces* was adjusted to 124.91 Mya (with the 95% Highest Probability Density (HPD) interval at 113.36 to 136.75 Mya) in the current analysis (Figure 5).

### AAI values

Inter-clade AAI values ranged from 69.07 to 72.03 % for GX, SR, and TM clades. Moderate AAI values were observed between the three novel clades and *Testudinimyces* (ranging from 69.38 ± 0.67 % for TM to 71.74 ± 0.79 % for GX). However, consistently low AAI values were observed between GX, SR, and TM clades and all mammalian AGF genera (52.65 to 66.21 %), and the fifth tortoise genus *Astrotestudinimyces* (60.26 to 63.69 %) (Table 4).

**Table 4:**
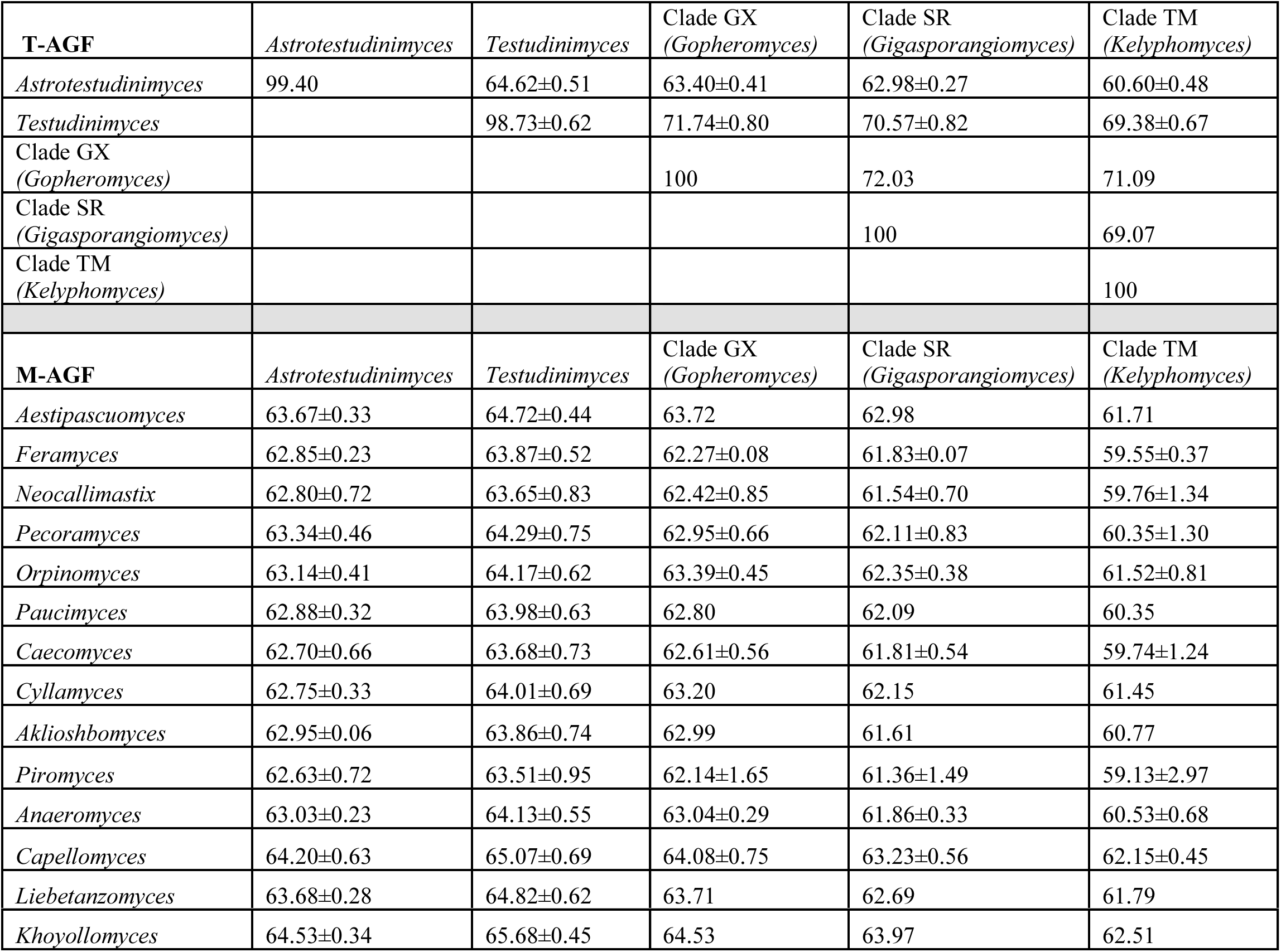
Genus-level amino acid identity (AAI) comparisons between tortoise-affiliated and mammal-affiliated AGF genera.

## Discussion

In this study, we report on the isolation of multiple anaerobic gut fungal (AGF) strains from tortoise feces. The isolates belonged to three different clades (clade GX from a Texas tortoise, and clades SR and TM from two different sulcata tortoises), bringing the total known tortoise-affiliated AGF (T-AGF) clades to five. Isolation efforts were initiated based on culture-independent surveys identifying samples harboring a high proportion of these novel clades (Table S2).^18^ Isolates GX1, GX2, and GX2A represent the first cultured representatives of genus NY56, a lineage identified in a previous culture-independent study.^18^ Collectively, the lack of studies on T-AGF (three, including this one), the low number of tortoise genera and species investigated so far (8 genera and 9 species out of 19 extant tortoise genera and 59 extant tortoise species), the relatively low number of tortoise subjects investigated so far (n = 11^18^ and two in this study), and the limited geographical distribution of examined subjects (all housed in zoos and wildlife reserves in the states of Oklahoma and Kansas, USA) could indicate that the scope of AGF diversity in tortoises might be broader than what is currently known. Additional efforts examining more tortoise taxa from other locations, as well as potential modifications to the isolation strategies, should expand T-AGF known diversity and enable culturing of additional novel T-AGF genera. Modification in the isolation protocols could be guided by results obtained here and in Pratt *et al.* (2023)^4^, e.g., using different substrates as well as adapting enrichment temperatures and incubation times. Phenotypically unique among AGF genera is the ability of clades SR and TM to grow on galactose, a substrate that is often recalcitrant to other AGF due to the absence of a galactokinase encoded in AGF genomes, and the ability of clade GX to grow on chitin.

Future comparative genomic efforts should explore the genetic basis behind these unique abilities. In addition to unique substrate preferences, the three clades exhibited individual differences in growth temperature ranges and optima, while enforcing broad patterns of differences between T-AGF and mammal-associated AGF (M-AGF). The observed temperature optima (32 to 35 °C in clade SR, and 35 °C, for clades GX and TM) and ranges (28 to 37°C for GX and SR clades, 22 to 37 °C for TM clade) are comparable to the other two genera previously isolated from tortoises (*Testudinimyces* (optimal, 30 °C; range, 30 to 39 °C), and *Astrotestudinimyces* (optimal, 35 to 39 °C; range, 30 to 39 °C)).^4^ These optima and ranges are different from the mammalian AGF genera, which have a narrower range of growth (usually 37 to 41 °C) with optimal growth at 39 °C (unpublished data).^4^ We argue that these lower optima/wider temperature ranges would aid in the survival and growth of T-AGF in their preferred poikilothermic (cold-blooded) hosts with lower and wider variation in internal temperature.^40^ Slower growth rates of some T-AGF taxa (e.g., clade GX) would mirror the slower basal metabolic rate of tortoises.^41^ In their hosts, long food retention time (12 to 14 days)^42^ would allow ample time for substrate colonization by slow-growing T-AGF.

Multiple lines of evidence (morphological as well as phylogenetic) support placing the obtained GX, SR, and TM clades into three distinct novel genera. Morphologically, the three clades exhibited unique microscopic features compared with closely related genera. The GX isolates showed abundant presence of elongated, thick, unbranched, nucleated structures on all media types investigated (Figure 1 D, G, J, L). Similar features have been previously observed in other AGF, like *Astrotestudinimyces divisus* (Figure 4s in Pratt *et al.*)^4^, *Testudinimyces gracilis* (Figure 3n in Pratt *et al.*)^4^, *Capellomyces elongatus* (Hanafy *et al*., Figure 5i)^43^, and *Ghazallomyces constrictus* (Figure 6j in Hanafy *et al.*)^43^, albeit not as frequently and consistently. In these previous publications the structures have been mentioned only briefly and were often identified as “elongated and irregular shaped sporangia”. In the GX isolates described in the present study, the structures likely were elongated sporangiophores (per definition, the stalk on which sporangia are developed) that failed to develop a terminal zoosporangium at their tip. Why this strain is producing these structures regularly and in large numbers remains unclear.

The SR isolates exhibited multiple thin hair-like filaments growing directly out of the sporangial walls (Figure 2I, J, L-P), a feature not previously observed in any other cultured AGF strain. These structures could be similar to the “zoospore-discharge tubes” previously reported for some *Chytridiomycota* like *Rhizophlyctis rosea* (Figure 2B-E in Letcher *et al.*)^44^, *Gorgonomyces chiangraiensis* (Figure 7K in Falcon *et al.*)^39^, or *Cladochytrium setigerum* (Figure 3C in Dubey and Upadhyay).^45^ In *Chytridiomycota*, however, these tubes are often wider than the filaments observed on SR0.6’s sporangia, and not as plentiful. No nuclei were identified in the hair-like filaments growing out of SR0.6 sporangia. Additionally, multiple sporangial pores were observed in SR isolates (Figure 2M, N), which appear to be the zoospore-release mechanism for this clade. To the best of our knowledge, zoospore release via multiple pores has not been observed previously in AGF or other, early diverging fungi.

The TM isolates showed fluidity in sporangial shape, a feature not previously observed in any other *Neocallimastigomycota* strain. While their sporangia started out circular, they kept growing irregularly, sometimes elongating in multiple directions (Figure 3D, I) and thinning at the tip, sometimes forming multiple circular protrusions (Figure 3K). While changes in sporangial morphology have been described in some AGF strains, these changes were linked to changes in carbon sources (e.g., *Liebetanzomyces*).^46^ The TM isolates, however, showed a wide variety of sporangial shapes on the same media in the same sample. This fluidity of sporangial shapes has, to the best of our knowledge, not yet been observed in early-diverging fungi.

Phylogenetic analysis using the phylogenetic markers ITS1, LSU, and RPB1 (Figure 4), as well as phylogenomic analysis using concatenated alignments of 758 single copy marker genes (Figure 5) placed the GX, SR, and TM isolates in three bootstrap supported clades, distinct from each other, as well as from the two other genera previously isolated from tortoises (*Testudinimyces* and *Astrotestudinimyces*). Sequence similarity thresholds to the closest relative, *Testudinimyces gracilis*, based on the D1-D2 region of LSU rRNA (Table 4), as well as inter-clade D1-D2 LSU rRNA similarity values (Table 4) were beyond those suggested for circumscribing genera (a minimum D1-D2 LSU sequence divergence threshold of 3 % from the closest cultured, validly described taxa).^2,32,47^ In addition, AAI values (ranging from 54.56 to 63.97%) to all other described AGF genera with a sequenced genome/transcriptome, exceed the previously suggested thresholds for circumscribing novel genera (AAI value of 85%) in the *Neocallimastigomycota*.^2,32,47^

Previous work using phylogenomics identified four statistically supported family-level clades in the *Neocallimastigomycota*^32^, with family boundaries based on phylogenomic tree topologies, as well as taxonomically informative morphological characteristics. The recent guidelines^32,47^ proposed AAI values of 75.0% as a guide for circumscribing families. AAI values of the five T-AGF clades compared to all M-AGF genera (average 63.16 ± 1.48%) are much lower than the suggested family cutoff of 75% (Table 5). Therefore, assignment of T-AGF to current *Neocallimastigomycota* families is unwarranted.

**Table 5:**
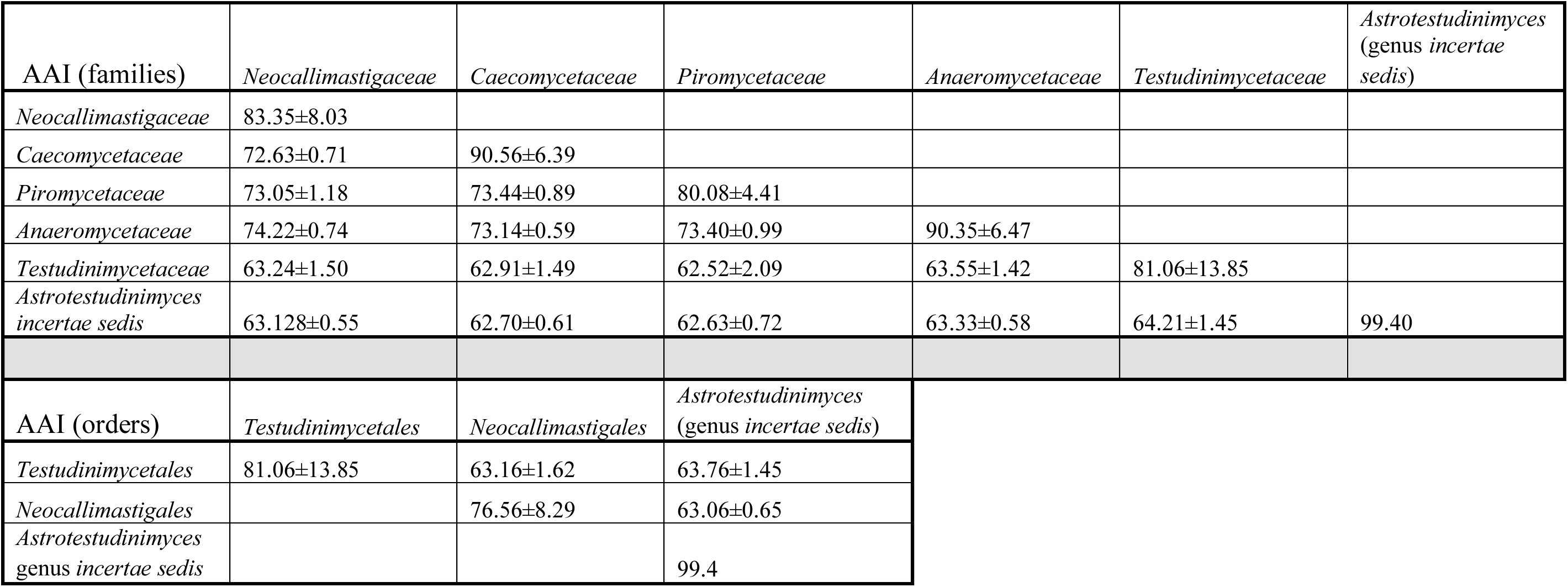
Family- and order-level amino acid identity (AAI) comparisons

Phylogenetic and phylogenomic analyses (Figures 4 and 5) show a clear branching between *Astrotestudinimyces* on one end, and *Testudinimyces* and the three novel genera described here on the other. This is also reflected in the lower AAI values of the *Testudinimyces*-affiliated genera to *Astrotestudinimyces* (64.21 ± 0.82%), which are again lower than the suggested family threshold of 75%. Hence, we propose accommodating four of the five T-AGF genera (*Testudinimyces* and the three novel genera described here) into a novel family (*Testudinimycetaceae,* named after the first described genus within the clade) showing average AAI values of 81.06 ± 13.9% amongst themselves. The solitary nature of *Astrotestudinimyces* renders accommodating it in a novel family unfeasible, since proposing novel families based on a single genus and species is unadvisable. As such, we propose designating *Astrotestudinimyces* as genus *incertae sedis* pending the description of additional closely related genera and species.

The current taxonomic outline of the phylum *Neocallimastigomycota* was proposed in 2023, prior to the discovery of T-AGF^4,32^, and encompasses one class (*Neocallimastigomycetes*), one order (*Neocallimastigales*), four families (*Neocallimastigaceae*, *Anaeromycetaceae*, *Piromycetaceae*, and *Caecomycetaceae*), and six genera *incertae sedis* (*Buwchfawromyces*, *Joblinomyces*, *Tahromyces*, *Agriosomyces*, *Aklioshbomyces,* and *Khoyollomyces*). T-AGF genera share a unique phylogenetic position in the *Neocallimastigomycota* phylogenomic tree (Figure 5). They are basal to all cultured M-AGF, share a unique ecological niche (tortoise gut)^18^, and display distinct phenotypic characteristics, Figures 1–3, Figure S3, S4, Table S3). This would justify proposing an intermediate rank (order) to accommodate some of the T-AGF genera and emend the description of the order *Neocallimastigales* to include only M-AGF genera. An AAI threshold for delineating orders in AGF has not been proposed before. Here, we argue that an AAI threshold of 65% is appropriate. AAI values between M-AGF genera and families (76.56 ± 8.29%) are higher than this cutoff, while AAI values between M-AGF and T-AGF (63.16 ± 1.62%) are lower than this threshold. Using that same threshold, genera in the family *Testudinimycetaceae* can be accommodated into a single order (average within order AAI values: 81.06 ± 13.85%), for which we propose the name *Testudinimycetales*. The genus *Astrotestudinimyces* has a unique position in the phylogenetic and phylogenomic tree (Figures 4 and 5) and shares lower AAI values with M-AGF (63.06 ± 0.65%) and with the genera in the proposed order *Testudinimycetales* (63.76 ± 1.45%, Table 4). As such, we propose keeping *Astrotestudinimyces* as genus *incertae sedis* pending the description of additional closely related genera and species. We further propose retaining all currently described AGF genera in a single class (*Neocallimastigomycetes*) in the phylum *Neocallimastigomycota*.

## Taxonomy

### Description of Gopheromyces

Alexandria Morris, Carrie J. Pratt, Mostafa S. Elshahed, Noha H. Youssef, Julia Vinzelj **gen. nov.** *MycoBank ID:* MB861080

*Typification*: *Gopheromyces tardescens* Alexandria Morris, Carrie J. Pratt, Mostafa S. Elshahed, Noha H. Youssef, Julia Vinzelj

*Etymology*: *Gopheromyces* (from *Gopherus*, the gopher tortoise, and *myces*, “fungus”) refers to host animal (*Gopherus berlandieri*) that the type species of this taxon was isolated from. An anaerobic filamentous fungus with polycentric thallus development, forming extensively branched nucleated hyphae and monoflagellated, ovoid or globose zoospores. Mature sporangia and free-swimming zoospores are rarely observed on rumen fluid cellobiose medium. The fungus produces characteristic elongated, unbranched, and nucleated stalks that might develop into zoosporangia under the right conditions. The clade is defined by the sequence deposited under BioProject PRJNA1345044; the type strain is *Gopheromyces tardescens* (strain GXA2).

### Description of Gopheromyces tardescens

Alexandria Morris, Carrie J. Pratt, Mostafa S. Elshahed, Noha H. Youssef, Julia Vinzelj **sp. nov.**

*MycoBank ID:* MB861081

*Typification*: Isolated in November 2024 from the frozen, then thawed feces of a Texas tortoise housed at the Oklahoma City Zoo (USA, Oklahoma, Oklahoma City, 35.52403791491748, -97.47250393174403). A metabolically active ex-type strain GX2 (that was isolated from the same fecal sample as the type strain GXA2) is maintained under anaerobic conditions and weekly transfer in rumen fluid cellobiose medium at 30 °C. It has been maintained at Oklahoma State University since its isolation in 2024. The holotype is stored at the Oklahoma State University (Stillwater, OK, USA) in 60% glycerol solution.

DNA and RNA extractions from the ex-type strain are stored at -80 °C at Oklahoma State University. The clade is defined by the sequence deposited under BioProject PRJNA1345044; the type strain is GXA2.

*Etymology*: ***Gopheromyces tardescens***. The epithet *tardescens* (Latin *tardescere*, “to grow slowly”) describes the slow growth rate observed in this taxon.

An obligate anaerobic filamentous fungus with polycentric thallus development and mature sporangia that are ovoid or bulb-shaped, with diameter 28 µm to 81 µm (average of 53.62 ± 28.14 µm). The fungus produces small globose monoflagellated zoospores averaging 8.11 ± 1.79 × 3.98 ± 0.68 µm (length x width). Produces thick, elongated, nucleated, unbranched structures with diameters of 8.15-36.24 µm (average of 21.26 ± 7.50 µm) in addition to wide and narrow hyphae. Mature sporangia and free-swimming zoospores are rarely observed on rumen fluid cellobiose medium but can be more frequent on medium containing pectin as sole carbon source. The type strain can survive on chitin as sole carbon source and can grow on a broad range of temperatures (28–37 °C, optimum: 35 °C). In liquid rumen fluid cellobiose media, the species produces white to slightly yellow biomass clusters that stick to the glass tubes but are detached when shaken. On RFC agar roll tubes, the type strain produces irregular shaped, white colonies that start out small and grow over time (up to 4.76 mm in diameter). The dense center darkens over time, while the fringed edges remain white to yellowish.

*Additional specimens examined*: Additional strains belonging to *Gopheromyces tardescens* were isolated from fecal samples of Texas tortoise (GX1 and GXA2), all of which were collected from: Oklahoma City Zoo, Oklahoma City, Oklahoma, USA, 35.52403791491748, -97.47250393174403).

### Description of Gigasporangiomyces

Taylor Mills, Samuel L. Miller, Mostafa S. Elshahed, Noha H. Youssef, Julia Vinzelj **gen. nov.**

*MycoBank ID:* MB861079

*Typification*: *Gigasporangiomyces pilosus*, Taylor Mills, Samuel L. Miller, Mostafa S. Elshahed, Noha H. Youssef, Julia Vinzelj.

*Etymology*: ***Gigasporangiomyces*** (from Greek *gigas*, “giant”, *sporangion*, “spore vessel”, and *myces*, “fungus”) refers to the large sporangia observed in the type species of this taxon. An anaerobic filamentous fungus with monocentric thallus development and monoflagellated zoospores. The fungus forms globose sporangia on short, slightly bulging sporangiophores. It is characterized by the formation of multiple thin filaments growing directly out of the sporangial wall and elongating outwards as the sporangia age. Mature sporangia can reach diameters of more than 100 µm. The clade is defined by the sequence deposited under BioProject PRJNA1345044; the type strain is *Gigasporangiomyces pilosus* (strain SR0.6).

### Description of Gigasporangiomyces pilosus

Taylor Mills, Samuel L. Miller, Mostafa S. Elshahed, Noha H. Youssef, Julia Vinzelj sp. nov.

*MycoBank ID:* MB861113

*Typification*: Isolated in March 2025 from the frozen, then thawed feces of a sulcata tortoise housed at the Tanganyika Wildlife Reserve (USA, Goddard, KS, 37.6727773364427, - 97.55778908935058). A metabolically active type strain, SR0.6, is maintained under anaerobic conditions and weekly transfer in rumen fluid cellobiose medium at 30 °C. It has been maintained at Oklahoma State University since its isolation. The holotype is stored at the Oklahoma State University (Stillwater, OK, USA) in 60% glycerol solution. DNA and RNA extractions from the type strain are stored at -80 °C at Oklahoma State University. The clade is defined by the sequence deposited under BioProject PRJNA1345044; the type strain is SR0.6.

*Etymology*: ***Gigasporangiomyces pilosus***. The epithet *pilosus* (from Latin, “hairy”) describes the thin, hair-like filaments growing out of the developing sporangia in this taxon.

An obligate anaerobic fungus with mature sporangia that are mostly globose (average diameters of 44.82 ± 21.10 µm, n = 50). Extraordinary large sporangia (average diameter of 139.99 ± 61.14 µm, n = 14) are regularly observed. The fungus produces small globose monoflagellated zoospores (5.79 ± 0.84 × 5.68 ± 0.88 µm = length x width, n = 15). Multiple thin, hair-like filaments (average width of 1.89 ± 0.25 µm, n = 25) are often growing directly out of the sporangial wall and elongate outwards as the sporangia age. Potential zoospore release mechanism through one or multiple pores forming in the sporangial wall. The type strain can grow on a broad range of temperatures (28–39 °C, optimum: 32–35 °C). In liquid rumen fluid cellobiose (RFC) media, the species produces white biomass flakes with denser centers that clump together over time to form irregularly shaped, loosely attached biomass clumps. On RFC agar roll tubes, the strain forms round colonies (3.2-4.8 mm in diameter), with white to yellowish filamentous edges that have a translucent shine to them.

*Additional specimens examined*: An additional strain belonging to *Gigasporangiomyces pilosus* was isolated from a fecal sample of sulcata tortoise (SR0.1), which was also collected from Tanganyika Wildlife Reserve (USA, Goddard, KS, 37.6727773364427, - 97.55778908935058).

### Description of Kelyphomyces

Taylor Mills, Samuel L. Miller, Mostafa S. Elshahed, Noha H. Youssef, Julia Vinzelj **gen. nov.**

*MycoBank ID:* MB861077

*Typification*: *Kelyphomyces adhaerens*. Taylor Mills, Samuel L. Miller, Mostafa S. Elshahed, Noha H. Youssef, Julia Vinzelj.

*Etymology*: ***Kelyphomyces*** (from Greek *kelyphos*, “shell, covering”, referring to the tortoise host *Centrochelys sulcata*) designates the tortoise-associated origin of the type species of this taxon.

An anaerobic filamentous fungus with polycentric thallus development and monoflagellated zoospores. The fungus is characterized by varying sporangia shapes and the formation of wide nucleated and narrow un-nucleated, but highly branched hyphae. Developing sporangia are globose at first, then grow out irregularly in all directions forming multiple different sporangial shapes. The clade is defined by the sequence deposited under BioProject PRJNA1345044; the type strain is TM0.3.

### Description of Kelyphomyces adherens

Taylor Mills, Samuel L. Miller, Mostafa S. Elshahed, Noha H. Youssef, Julia Vinzelj **gen. nov.**

*MycoBank ID:* MB 861078

*Typification*: Isolated in April 2025 from the frozen, then thawed faeces of a sulcata tortoise housed at Amarillo Zoo (USA, Amarillo, OK, 35.239593462795916, -101.83516564339884). A metabolically active type strain TM0.3 is maintained under anaerobic conditions and weekly transfer in rumen fluid cellobiose medium at 30 °C. It has been maintained at Oklahoma State University since its isolation. The holotype is stored at the Oklahoma State University (Stillwater, OK, USA) in 60% glycerol solution. DNA and RNA extractions from the type strain are stored at -80 °C at Oklahoma State University. The clade is defined by the sequence deposited under BioProject PRJNA1345044; the type strain is TM0.3. *Etymology*: ***Kelyphomyces adherens***. The epithet *adhaerens* (Latin = “sticking, clinging”), describes the strong adherence of this fungus to the glass surface in liquid media.

An anaerobic filamentous fungus with polycentric thallus development and monoflagellated zoospores. The fungus is characterized by varying sporangia shapes and the formation of wide nucleated and narrow un-nucleated, but highly branched hyphae. Developing sporangia are globose at first (average length x width = 16.84 ± 3.79 × 19.37 ± 4.47 µm, n = 8), then grow out irregularly in all directions forming multiple different sporangial shapes. The fungus produces small globose monoflagellated (rarely biflagellated) zoospores (average length x width = 5.03 ± 0.55 × 5.25 ± 0.60 µm, n = 21). The species can grow at 22 to 37 °C (optimum, 35 °C). In liquid rumen fluid cellobiose (RFC) media, the strain forms tight, small biomass clusters that quickly (within a day) turn brown and adhere so tightly to the glass tube it must be scraped off. On RFC agar rolltubes colonies are small, with dense dark centers and filaments growing out of them in all directions.

Description of order *Testudinimycetales*, ord. nov.

Obligate anaerobic fungi that produces monoflagellated zoospores, mono- or poly-centric thalli and a filamentous rhizoidal system. The clade is circumscribed by phylogenomic analysis and AAI values, and confirmed by LSU and RPB1 phylogenetic analyses, as well as morphological characteristics. It currently accommodates the family *Testudinimycetaceae*.

*Type family: Testudinimycetaceae MycoBank ID:* MB861075

Description of Family *Testudinimycetaceae,* fam. nov.

Obligate anaerobic fungi that produce monoflagellated zoospores, mono- or polycentric thalli and a filamentous rhizoidal system. The clade is circumscribed by phylogenomic analysis and AAI values, and confirmed by LSU and RPB1 phylogenetic analyses, as well as morphological characteristics. Currently accommodates the genera *Testudinimyces, Gopheromyces, Gigasporangiomyces, and Kelyphomyces*.

*Type genus*: *Tesudinimyces MycoBank ID:* MB861076

#### Emended description of order *Neocallimastigales*

Obligate anaerobic fungi that produce mono- or poly-flagellated zoospores, mono- or poly-centric thalli and a filamentous or bulbous rhizoidal system. The clade is circumscribed by phylogenomic analysis and AAI values, and confirmed by LSU and RPB1 phylogenetic analyses, as well as morphological characteristics. Currently accommodates the families *Neocallimastigaceae*, *Anaeromycetaceae*, *Piromycetaceae*, and *Caecomycetaceae*, and the 6 *incertae sedis* genera (*Buwchfawromyces*, *Joblinomyces*, *Tahromyces*, *Agriosomyces*, *Aklioshbomyces,* and *Khoyollomyces).* The emended description of the order *Neocallimastigales* is generally similar to that previously provided for the order (https://cdnsciencepub.com/doi/10.1139/b93-044), with amendments to exclude the tortoise-affiliated genera *Testudinimyces*, *Gopheromyces*, *Gigasporangiomyces*, and *Kelyphomyces*, and to circumscribe its boundaries to encompass a monophyletic clade of four families and six genera *incertae sedis*, all isolated from mammalian sources.

*Type family: Neocallimastigaceae MycoBank ID:* MB 90438 **Genera *incertae sedis***

The following genus is assigned as genera *incertae sedis* within the class *Neocallimastigomycetes*: *Astrotestudinimyces* [19] MycoBank ID: 847431.

## Data availability

Sequencing data generated in this study is available through NCBI (BioProject number PRJNA1345044). Data pertaining to the physiology of the described taxa are published in this manuscript and accompanying supplementary material. More information on the strains can be obtained from the authors upon request.

## Acknowledgements

We would like to thank the Oklahoma City Zoo, the Amarillo City Zoo, and the Tanganyika Wildlife Park for their contribution of samples and the smooth collaboration. We would further like to thank Stephen Marek for sharing his insights on fungal morphology and the Boren Veterinary Medical Teaching Hospital for their continuous support in providing rumen fluid.

## Author Contributions

Conceptualization – MSE, NHY. Data curation – JV, NHY. Formal analysis – JV, YW, NHY. Funding acquisition – MSE, NHY, YW. Investigation – AM, TM, CP, JV, NHY. Project administration – JV, MSE, NHY. Resources – MSE, NHY, SLM, JV. Supervision – JV, MSE, NHY. Visualization – AM, JV, NHY. Writing original draft – JV, NHY, MSE. Writing review/edition – all authors.

