## Supplementary Figures for "Description of *Gopheromyces tardescens*, gen. nov., sp.nov., *Gigasporangiomyces pilosus,* gen. nov., sp.nov., *Kelyphomyces adhaerens,* gen. nov., sp. nov., proposal of *Testudinimycetales* ord. nov. and *Testudinimycetaceae* fam. nov., and emended description of the order *Neocallimastigales*"

**Keywords:** *Neocallimastigomycota*; anaerobic fungi; tortoises; AAI; molecular dating analysis

**Figure S1. Growth curves for the three novel *Neocallimastigomycota* isolates (GXA2, SR0.6, TM0.3) grown in rumen fluid media containing either cellobiose (SR0.6, TM0.3) or lactose (GXA2) as substrate at 35 °C.** Solid lines represent the average cumulative gas pressure (PSI) and dashed lines the average of the visual growth evaluation (from 0, no growth, to 4, very good growth). Measurements were taken from the same four tubes per strain at the indicated time points after subculture.

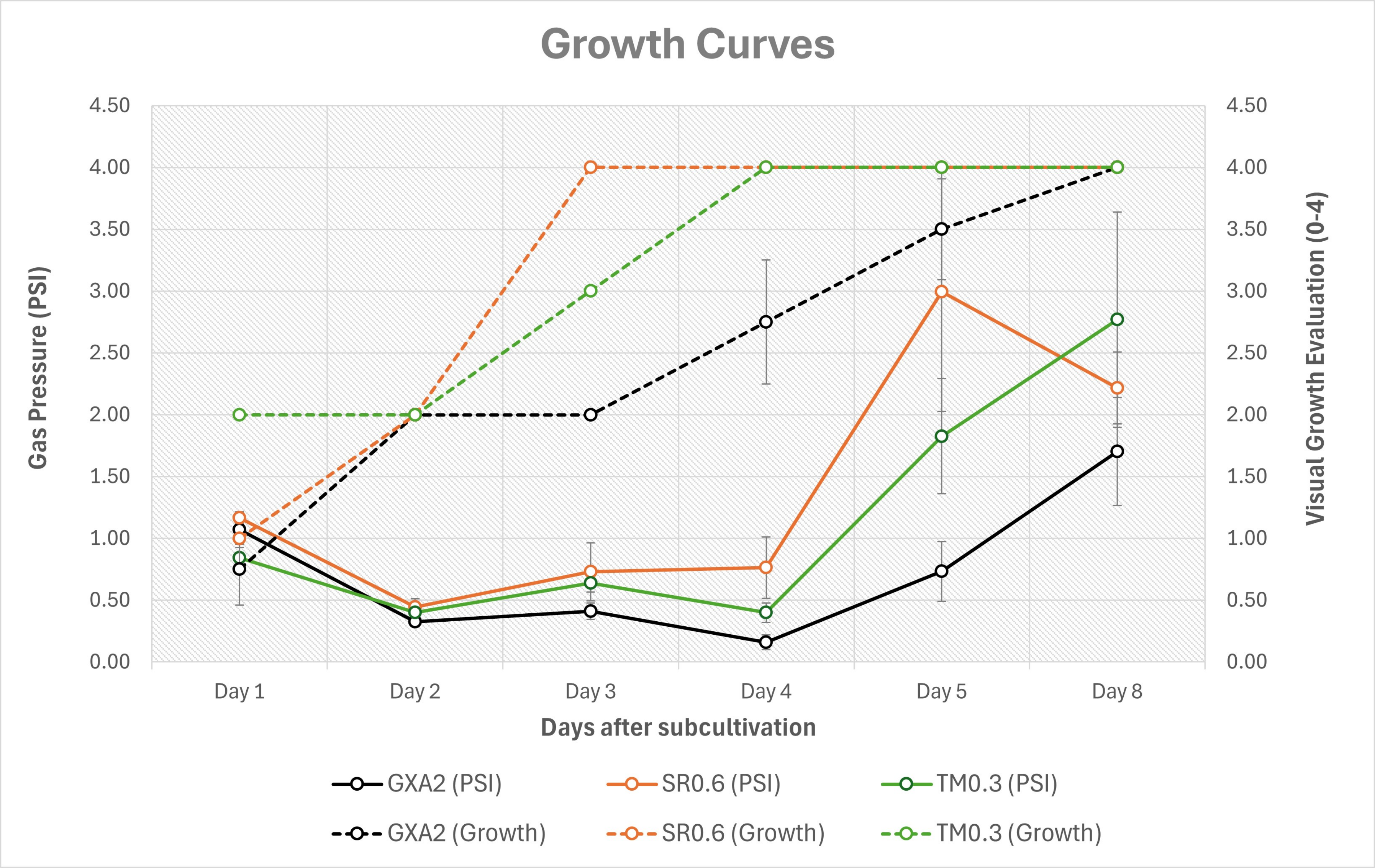
Growth Curves
Gas Pressure (PSI)
Visual Growth Evaluation (0-4)
4.50
4.00
3.50
3.00
2.50
2.00
1.50
1.00
0.50
0.00
Day 1
Day 2
Day 3
Day 4
Day 5
Day 8
Days after subcultivation
GXA2 (PSI)
SR0.6 (PSI)
TM0.3 (PSI)
GXA2 (Growth)
SR0.6 (Growth)
TM0.3 (Growth)

**Figure S2. Morphology of strain GXA2 (Clade GX, *Gopheromyces tardescens*), additional images.**

**(A)** Strain GXA2 grown on rumen fluid media containing 0.1% w/v switchgrass as carbon source. **(B-J)** Strain GXA2 grown on rumen fluid media containing cellobiose as carbon source. **(A-C)** SEM microscopy, **(D, E, H-J)** Phase contrast microscopy, **(F, G)** Confocal microscopy with DAPI. Images show the extensive hyphal entanglement exhibited by GXA2 **(A, C)**, the sporangia produced **(E, F, I, J)**, and the elongated stalks characteristic of GXA2 **(B, D, H)**. Scale bars indicate 100 µm **(A-C)**, 50 µm **(F, G)**, 25 µm **(D, E, H-J)**.

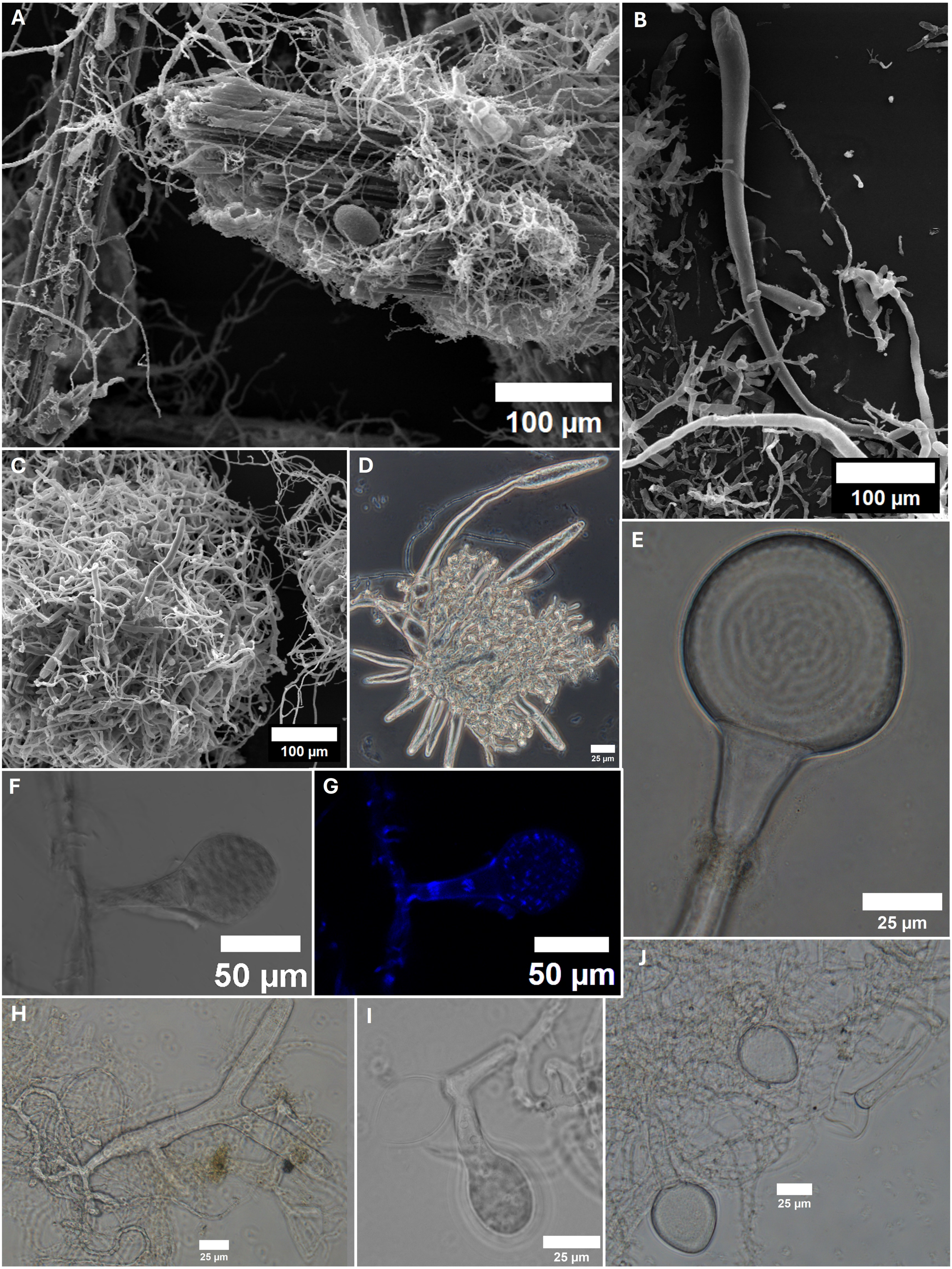

A
100 μm
B
100 μm
C
100 μm
D
25 μm
E
25 μm
F
50 μm
G
50 μm
H
25 μm
I
25 μm
J
25 μm

**Figure S3. Morphology of strain SR0.6 (Clade SR, *Gigasporangiomyces pilosus*), additional images.**

**(A-F)** Strain SR0.6 grown on rumen fluid media containing cellobiose as carbon source. **(A, C-F)** Phase contrast microscopy. **(B)** Confocal microscopy shows a large, ovoid sporangia entangled in filaments (especially at the base, black arrow). **(C/D)** DAPI staining showing monocentric thalli (nuclei only in the sporangia) and the thin, hair-like filaments growing out of the sporangial walls (black arrows). Autofluorescence shows the hair-like filaments as well as the sporangiophores more clearly (white arrows). Scale bars indicate 50 µm **(A-D, F)** or 20 µm **(E)**.

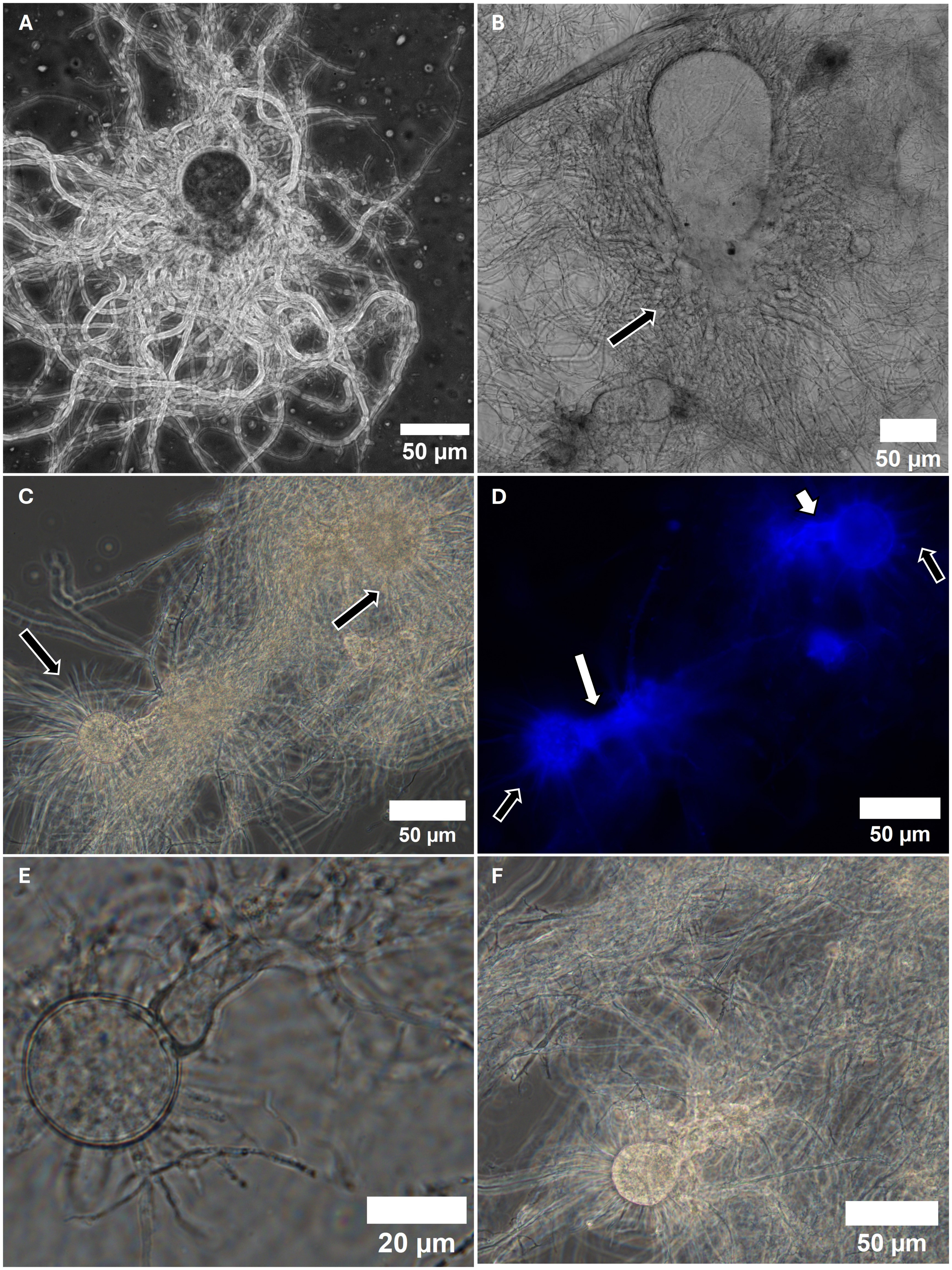
A
50 µm
B
50 µm
C
50 µm
D
50 µm
E
20 µm
F
50 µm

**Figure S4. Morphology of strain TM0.3 (Clade TM, *Kelyphomyces adhaerens*), additional images.**

Phase contrast microscopy of strain TM0.3 grown on rumen fluid media containing cellobiose **(A-E)** or inulin **(D-H)** as carbon source showing more of the various sporangial shapes observed. Scale bars indicate 50 µm **(A, C, E, F, G)** or 20 µm **(B, D, H)**.

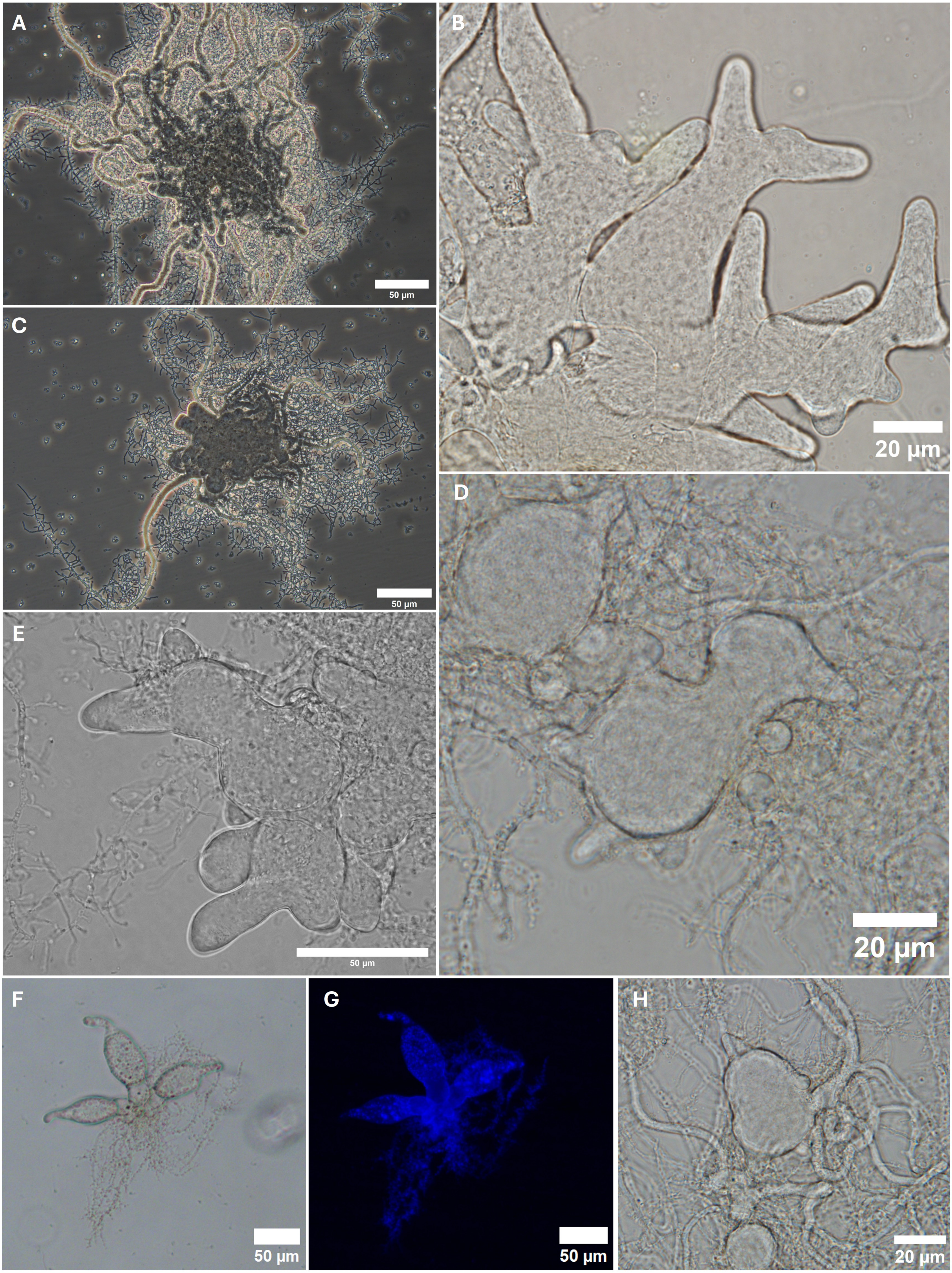
A
50 µm
B
20 µm
C
50 µm
D
20 µm
E
50 µm
F
50 µm
G
50 µm
H
20 µm

**Figure S5. Growth of the three novel *Neocallimastigomycota* isolates (GXA2, SR0.6, TM0.3) in rumen fluid media containing various substrates.** Bars represent the average gas pressure (PSI), shorter horizontal lines represent the average of the visual growth evaluation (from 0, no growth, to 4, very good growth), and the black horizontal lines represent the average gas pressure of the negative controls (uninoculated tubes). **(A)** Results after the first transfer on the various substrates. **(B)** Average after the last transfer on the various substrates. **(C)** Average over the whole experiment.

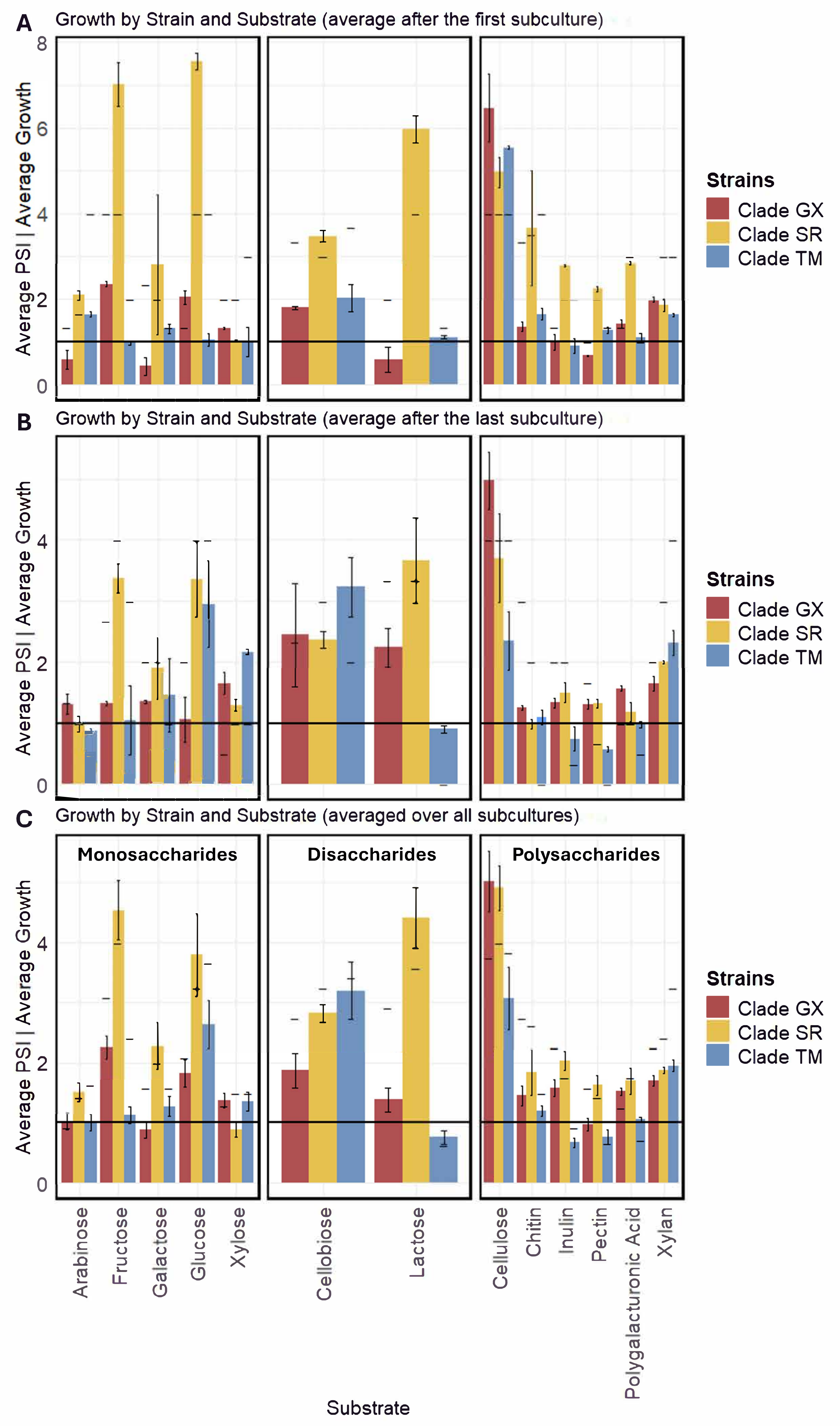
A
Growth by Strain and Substrate (average after the first subculture)
B
Growth by Strain and Substrate (average after the last subculture)
C
Growth by Strain and Substrate (averaged over all subcultures)
Monosaccharides
Disaccharides
Polysaccharides
Average PSI | Average Growth
Strains
Clade GX
Clade SR
Clade TM
Arabinose
Fructose
Galactose
Glucose
Xylose
Cellobiose
Lactose
Cellulose
Chitin
Inulin
Pectin
Polygalacturonic Acid
Xylan
Substrate

**Figure S6. Growth of the three novel *Neocallimastigomycota* isolates (GXA2, SR0.6, TM0.3) at various temperature. Strains were either grown in rumen fluid medium containing cellobiose (SR0.6, TM0.3) or lactose (GXA2) at different temperatures.** Bars represent the average gas pressure (PSI), shorter horizontal lines represent the average of the visual growth evaluation (from 0, no growth, to 4, very good growth), and the black horizontal lines represent the average gas pressure of the negative controls (uninoculated tubes). **(A)** Results after the first transfer at the various temperatures. **(B)** Average after the last transfer at the various temperatures. For strain GXA2 on 22 and 39 °C the data after the second subculture is displayed. **(C)** Average over the whole experiment. For strain GXA2 on 22 and 39 °C the data was averaged just over the first and second subculture since it was dead and not further subcultured.

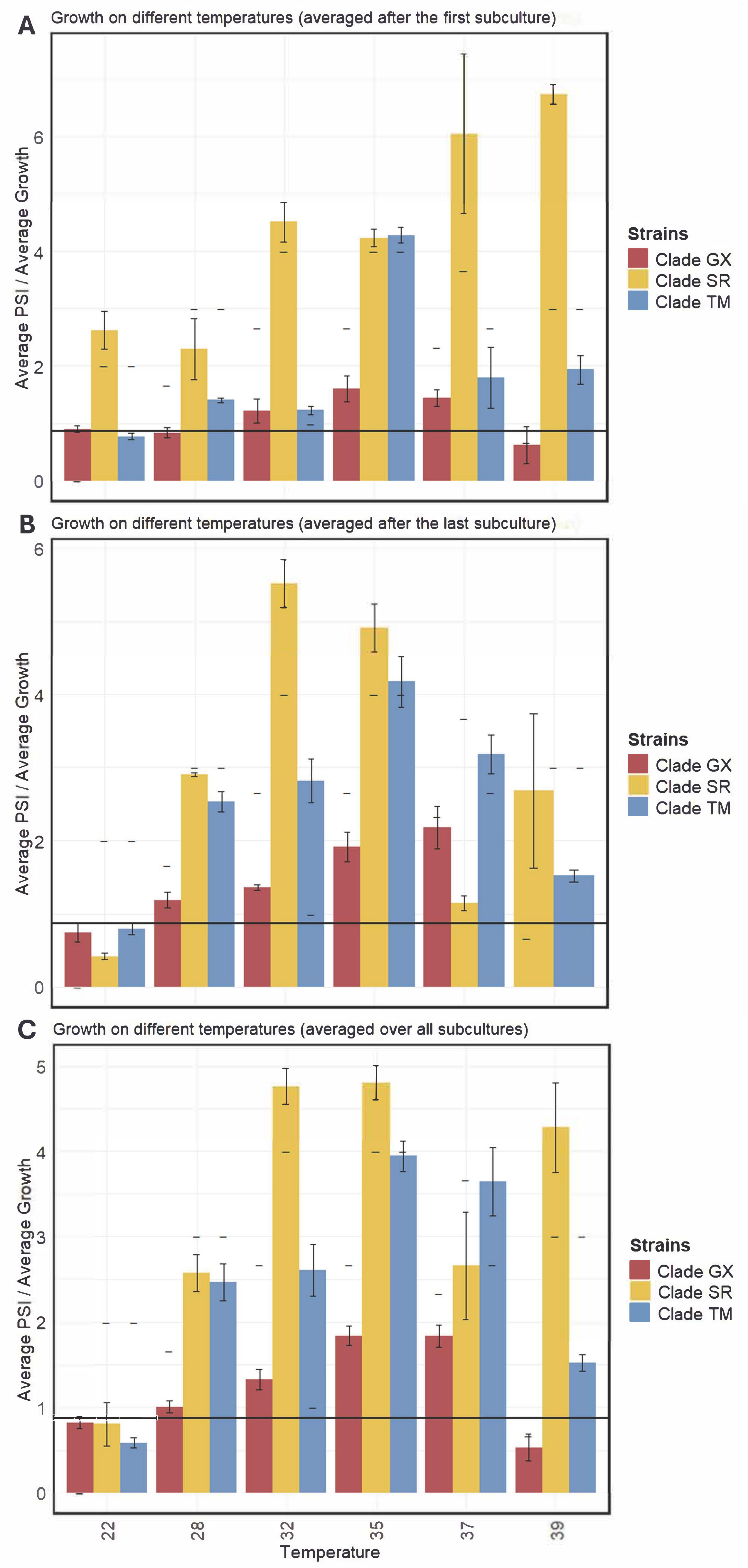
A
Growth on different temperatures (averaged after the first subculture)
B
Growth on different temperatures (averaged after the last subculture)
C
Growth on different temperatures (averaged over all subcultures)
Average PSI / Average Growth
Temperature
22
28
32
35
37
39
Strains
Clade GX
Clade SR
Clade TM
